# A bacterial ribosome hibernation factor with evolutionary connections to eukaryotic protein synthesis

**DOI:** 10.1101/2022.11.24.517861

**Authors:** Karla Helena-Bueno, Chinenye L. Ekemezie, Charlotte R. Brown, Arnaud Baslé, James N. Blaza, Chris H. Hill, Sergey V. Melnikov

## Abstract

During starvation and stress, virtually all organisms arrest protein synthesis to conserve energy. Inactive ribosomes are converted into a dormant state, in which they are protected from damage by hibernation factor proteins. In bacteria, two major families of hibernation factors have been described, but the low conservation of these proteins and the huge diversity of species, habitats, and environmental stressors has confounded their discovery. In this study, using proteomics and cryo-EM, we identify a new dormancy factor from the psychrophilic bacterium *Psychrobacter urativorans*. By isolating ribosomes under cold-shock conditions, we observe a previously unknown protein bound to the ribosomal A site, protecting critical elements of both the decoding and peptidyl transferase centers. We show that this new factor, which we term Balon, is a homolog of the archaeo-eukaryotic translation factor aeRF1, providing a long-predicted evolutionary “missing link” between the eukaryotic and bacterial translation machinery. Our structures reveal that Balon is delivered to both vacant and actively translating ribosomes by EF-Tu, highlighting an unexpected and previously unknown role for this elongation factor in the bacterial stress response. We describe several unique structural motifs that allow Balon to bind ribosomes in an mRNA-independent manner, initiating a new mode of ribosome dormancy that can commence while ribosomes are still engaged in protein synthesis. Our bioinformatic analysis shows that putative Balon-encoding genes can be found within stress-response operons in nearly 20 % of all known bacterial species, including many human pathogens. Taken together, our work suggests that Balon/EF-Tu regulated ribosome dormancy is likely to be a ubiquitous stress-response mechanism throughout the bacterial kingdom. These findings call for a revision of our model of bacterial translation inferred from common model organisms and hold numerous implications for how we understand and study ribosome dormancy.

## Introduction

When a living cell encounters environmental stress, metabolic activity is greatly reduced until conditions improve. Until recently, this was believed to be a passive process in which enzymes simply become idle, with vacant active sites. It is now clear that all organisms—from bacteria to humans—employ specific mechanisms to protect critical cellular machinery from damage during stress. This involves placing key enzymes and macromolecular assemblies into a controlled state of molecular hibernation^1,2^.

The phenomenon of molecular hibernation has been observed for many cellular enzymes, including RNA polymerases^3-5^, proteasomes^6^, and catalases^7^. Most extensively, this process has been studied in ribosomes – essential ribonucleoprotein complexes that catalyse protein synthesis, also known as translation. During normal conditions, ribosomes bind their ligands, such as mRNA and tRNAs, to perform protein synthesis^8-11^. However, during starvation and stress, ribosomes terminate protein synthesis, dissociate from mRNA and tRNAs, and enter molecular hibernation by associating with a specialized group of proteins known as hibernation factors or dormancy factors^12,13^.

Initial studies of ribosome hibernation factors suggested they may arrest protein synthesis due to their occupation of ribosomal binding sites for mRNA and tRNAs^14,15^. However, recent biochemical and genetic studies have revealed that dormancy factors also protect ribosomes from degradation by shielding their vulnerable active centres from cleavage by cellular nucleases^16-18^. Therefore, when ribosome hibernation factors are depleted through genetic knockouts, cells do not lose their ability to arrest protein synthesis in response to stress; rather, these cells rapidly accumulate defective ribosomes, causing a slower or incomplete recovery from stress. Hibernation factors thereby endow cells with a competitive advantage: the preservation of ribosomes during starvation, stress or drug treatment enables them to rapidly resume growth when the environment becomes more favourable^19-22^.

Ribosome dormancy factors have been found in some species, yet it is unknown how many exist in nature. To date, several families of ribosome dormancy factors have been characterized in eukaryotes, including Stm1 in yeast^23^, Serpbp1 and Irfd2 in mammals^24,25^, Lso2 in mammals and fungi^26-28^, and Mdf1 in the fungal parasites microsporidia^29,30^. Two major families of dormancy factors have also been described in bacteria, including the RaiA family (including RaiA, Hpf and LrtA), found in many bacterial lineages, and Rmf, found in some γ-proteobacteria^12,15,31,32^. Unlike most core translation factors, dormancy factors are highly diverse and lack conservation even within a single domain of life, thus complicating their discovery.

Structural analyses of dormant ribosomes from several species have revealed an array of structurally dissimilar proteins that bind to different ribosomal sites^15,23-30^. Nonetheless, all dormancy factors share two common properties: they bind almost all cellular ribosomes during conditions of starvation or stress, and they protect the most critically important functional sites in the ribosome, such as the mRNA channel and tRNA-binding sites^12^. However, the lack of universal conservation of ribosome dormancy factors, along with a small number of suitable organisms to study ribosome dormancy, leaves it unclear to what extent findings made in common model organisms—mainly *E. coli*—can be considered generally representative.

To address this, here we introduce a new model organism: the cold-adapted bacterium *Psychrobacter urativorans*. By examining *P. urativorans* ribosomes isolated under conditions of cold shock, we discover a new mechanism of translational response to stress. Using cryo-EM, we observe that *P. urativorans* ribosomes enter a state in which the A site is occupied by an uncharacterized protein that shields two active centres of the ribosome: the decoding centre and the peptidyl-transferase centre. Strikingly, in contrast to previously identified hibernation factors, this factor engages not only with vacant ribosomes but also with ribosomes associated with mRNA and tRNAs.

Homologs of this new factor are present in nearly 20 % of bacterial genomes, though notably absent from common model organisms such as *E. coli, S. aureus*, and *B. subtilis* – explaining why it has been undetected until now and emphasizing the importance of venturing beyond typical mesophilic bacteria to discover new biology. Our cryo-EM analysis indicates that this newly identified protein is a distant homolog of the eukaryotic translation factors eRF1 and Pelota that participate in other aspects of the translation process, not dormancy. We therefore name this protein Balon (Spanish for “Ball”) due to its distant structural and sequence similarity to Pelota (also Spanish for “Ball”). Our study provides the first experimental evidence of the existence of eukaryote-type translation factors in bacterial cells, illuminating a long-sought evolutionary connection between the bacterial and eukaryotic protein synthesis machinery^33-35^. Overall, our discovery and characterization of Balon demonstrates that bacteria can employ a qualitatively distinct and previously unknown mechanism of translational stress response compared to the generalized model of ribosome hibernation based on studies in *E. coli*, offering broad implications for how we understand and study the process of ribosome dormancy in response to stress.

## Results

### Cryo-EM analysis of ribosomes from ice-treated bacteria identifies a new factor of ribosome dormancy, Balon

Our interest in ribosome hibernation arose from our studies of protein synthesis at cold temperatures, using the bacterium *P. urativorans* as a model organism. *P. urativorans* is a notorious cold-adapted organism that can spoil frozen food due to its ability to grow at sub-zero temperatures^36-39^. In the laboratory, *P. urativorans* has a recommended growth temperature of 10 °C, but in nature *P. urativorans* can survive much colder environments, including Antarctic permafrost soil and glacial ice^36^.

To understand how *P. urativorans* adapts protein synthesis to sudden decreases in temperature, we isolated ribosomes for analysis shortly after inducing cold shock. As *P. urativorans* are slow-growing bacteria, with a doubling time greater than one day, we first grew cultures for several days at 10 °C to produce sufficient biomass. We then induced cold shock by exposing these growing cultures to ice for 30 minutes before extracting ribosomes for proteomic and cryo-EM analyses (**Methods, Fig. S1**). Previous studies have shown that in organisms such as *E. coli* and *B. subtilis*, sudden decreases in temperature lead to formation of mRNA secondary structures that result in an immediate and nearly complete halt of protein synthesis for several hours until cells produce high quantities of the stress-response protein CspA (up to 12% of the total protein content), which unfolds mRNAs and enables translation at colder temperatures^40-42^. We therefore anticipated a similar arrest of protein synthesis in *P. urativorans* in response to ice treatment.

Consistently, we found that cold shock of *P. urativorans* cells leads to a rapid depletion of polysomes and accumulation of monosomes, indicating that ribosomes disengage from protein synthesis (**Fig. 1A**). Our mass spectrometry analysis revealed that after 30 minutes of ice treatment, there was only a slight increase in the CspA content in *P. urativorans* cells (1.3 fold), indicating that the cells are in the early stages of a cold shock response (**Supplementary Data S1**). Our subsequent cryo-EM analysis revealed that, in contrast to actively translating ribosomes that exist in multiple conformations, nearly all the ribosomes isolated from the ice-treated cells adopt a uniform, non-rotated conformation. Furthermore, the A site of these ribosomes is no longer bound to canonical factors of protein synthesis such as aminoacyl-tRNAs or the elongation factor EF-G. Instead, in the majority of our dataset (∼66 %) the A site is occupied by a previously uncharacterized protein, which we term Balon (**Figs. S2-S5**).

**Fig 1.**
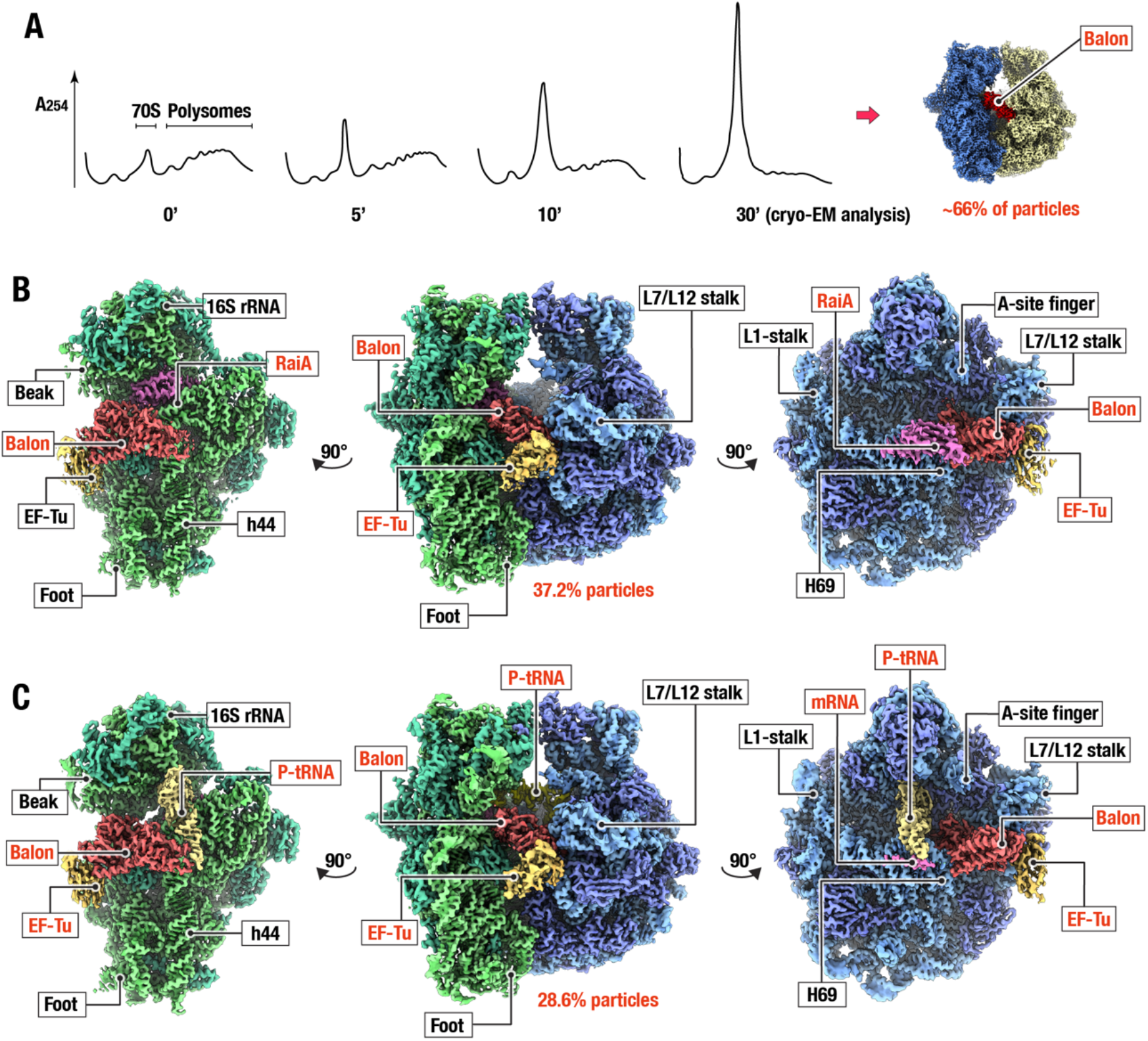
Cryo-EM analysis of ribosomes from ice-treated bacteria identifies a new factor of ribosome dormancy, Balon. (**A**) Polysome profiling in sucrose gradients shows an accumulation of monomeric ribosomes during the first 30 minutes of response to ice treatment of *P. urativorans* cells. (**B, C**) Cryo-EM maps at 3.1 Å resolution depict the two most prevalent states of ribosomes isolated from bacteria *P. urativorans* during cold shock. Each density map is colour-coded and shown in three orthogonal views. The 30S subunit (green) is shown alone with bound factors on the left, the 50S subunit (blue) is shown alone with bound factors on the right, and the full 70S particle is shown in the middle panel. (**B**) The first state corresponds to ∼37 % of ribosomes in the sample and consists of ribosomes bound to a previously unknown translation factor, Balon, and the known hibernation factor RaiA. (**C**) The second state corresponds to ∼29 % of ribosomes in the sample. This state represents ribosomes bound to Balon, mRNA, and P-site tRNA. Both states of the ribosome also show the presence of the elongation factor EF-Tu bound to Balon.

Of the Balon-bound ribosomes, particle classification revealed that approximately half (∼ 57 %) are also bound to the ribosome hibernation factor RaiA, and another half (∼ 43 %) do not contain RaiA, but are bound to heterologous mRNA and peptidyl-tRNA in the P/P state (**Fig. 1 B,C**). The cryo-EM data, along with our mass-spectrometry analysis of the crude ribosome samples (**Supplementary Data S2**) indicate that Balon-containing ribosomes are also associated with the elongation factor EF-Tu, which binds Balon in a similar fashion to aminoacyl-tRNAs when it delivers them to the ribosomal A site during protein synthesis (**Fig. 1 B,C, Fig. S6**). Overall, our structural analysis reveals that *P. urativorans* cells contain an additional protein that exhibits two common traits of all previously identified ribosome hibernation factors: it binds a large proportion of cellular ribosomes under stress conditions, and it occupies the ribosomal active site, making it physically inaccessible to other molecules.

### Balon resembles archaeal and eukaryotic protein synthesis factors

To reveal the identity and better understand the evolutionary origin and molecular function of Balon, we assessed whether Balon structurally resembles any known protein family. Using our cryo-EM map, we first built a poly-alanine model of Balon, and then used this model to search for structurally similar proteins across the Protein Data Bank (**Methods**). Interestingly, despite Balon being a bacterial protein, our search revealed greatest structural similarity to the aeRF-1 family of proteins from archaea and eukaryotes (**Fig. 2A, Table S2**). This family includes two ribosome-binding proteins: (1) aeRF1, a translation termination factor that binds the ribosomal A site and terminates protein synthesis in response to mRNA stop codons, and (2) Pelota, a ribosome-rescue factor that binds the A site on ribosomes that are arrested by aberrant mRNAs^22,43,44^. Our subsequent mass spectrometry analysis of the crude *P. urativorans* ribosome samples revealed the identity and sequence of Balon—a 41kDa protein, currently annotated as hypothetical protein AOC03_06830 (**Supplementary Data 2**).

**Fig 2.**
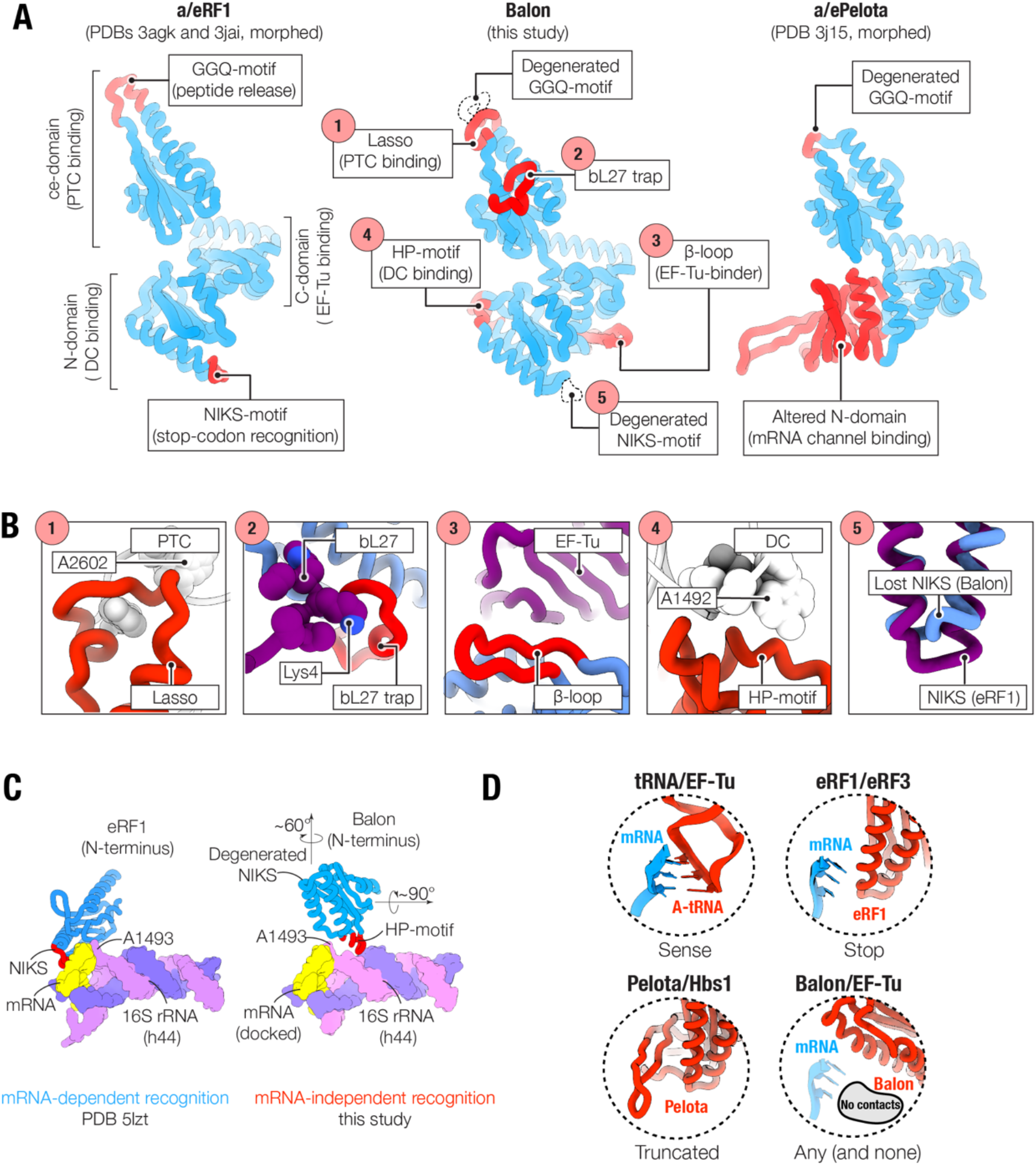
Balon resembles proteins aeRF1 and Pelota, indicating a shared evolutionary origin with the aeRF1-family of archaeo-eukaryotic translation factors. (**A**) Comparison illustrating structural similarity between Balon, aeRF1, and Pelota proteins. The overall structure of Balon most closely resembles the translation termination factor aeRF1: it shares a common domain organization and binds the ribosome in a similar way via contacts with EF-Tu, the decoding centre (DC) and the peptidyl-transferase centre (PTC). Balon lacks key structural features that are required for aeRF1 activity, including the NIKS-motif (stop-codon recognition) and the GGQ-motif (nascent peptide release). (**B**) Five unique structural features (labelled) distinguish Balon from both aeRF1 and Pelota and allow Balon to bind dormant bacterial ribosomes. (**C**) Comparison illustrating the dissimilar mechanisms by which eaRF1 and Balon (blue) recognize the decoding centre of the ribosome (16S rRNA and mRNA fragment shown for simplicity). (**D**) Comparison of decoding site-binding strategies used by Balon and other ligands of the ribosomal A site.

A structure-guided sequence alignment showed that Balon shares just ∼ 10 % pairwise sequence identity with both aeRF1 and Pelota, yet these three proteins have a similar overall architecture (rmsd < 4.0 Å for their structurally conserved core comprising approximately 280 of 400 Cα atoms). Furthermore, these three proteins have a similar length of ∼ 370 (Balon), ∼ 400 (aRF1) and ∼ 410 amino acids (Pelota) and the same domain organization, including the N-terminal domain that binds the small ribosomal subunit, the central domain that binds the large ribosomal subunit, and the C-terminal domain that binds conserved translation factor EF-Tu and its homologs in eukaryotes and archaea (**Fig. 2A-D, Fig. S6-S8**). This structural similarity suggests a common evolutionary origin for Balon, aeRF1 and Pelota and demonstrates that bacteria possess a functional translation factor from the aeRF1-family. Importantly, this represents the first experimental evidence of functional eukaryotic-like translation factors in bacteria.

### EF-Tu can act as a stress-response factor that delivers Balon to the ribosomal active site

Our cryo-EM maps and mass-spectrometry analysis of crude ribosome samples suggested that Balon may bind to the ribosome in complex with the universally conserved factor of protein synthesis EF-Tu (**Fig. 1B, Supplementary Data S2**). However, our first cryo-EM dataset showed only very weak density corresponding to EF-Tu, likely due to EF-Tu dissociation from the ribosome particles, which commonly happens after EF-Tu delivers substrates to the ribosomal A site^45^. We therefore collected an additional cryo-EM dataset and observed a large increase in the number of particles with bound EF-Tu (**Fig. S5, Methods**). This allowed us to obtain a better map that revealed the overall structure of EF-Tu bound to Balon in the ribosomal A site, uncovering the details of this molecular interaction (**Fig. 3A, Fig. S5, Methods**). Our analysis revealed that Balon binds the C-terminal domain of the EF-Tu molecule, similar to aeRF1 and Pelota binding the C-terminal domains of EF-Tu homologs known as eRF3 and Hbs1, respectively (**Fig. 3B, C**). This binding similarity can explain the simultaneous presence of EF-Tu and Balon in our ribosome samples: in archaea, EF-Tu delivers either aeRF1 or Pelota to the ribosomal A site to terminate translation or reactivate arrested ribosomes, respectively^46^, suggesting a conserved mechanism of delivery for aeRF1, Pelota, and Balon to the ribosome by EF-Tu (**Fig. 3C**). Importantly, this observation reveals a previously unknown biological activity of the elongation factor EF-Tu. This factor was believed to remain inactive during stress, however here we observe that it facilitates the delivery of a stress-response protein to the ribosome (**Fig. 3D**).

**Fig 3.**
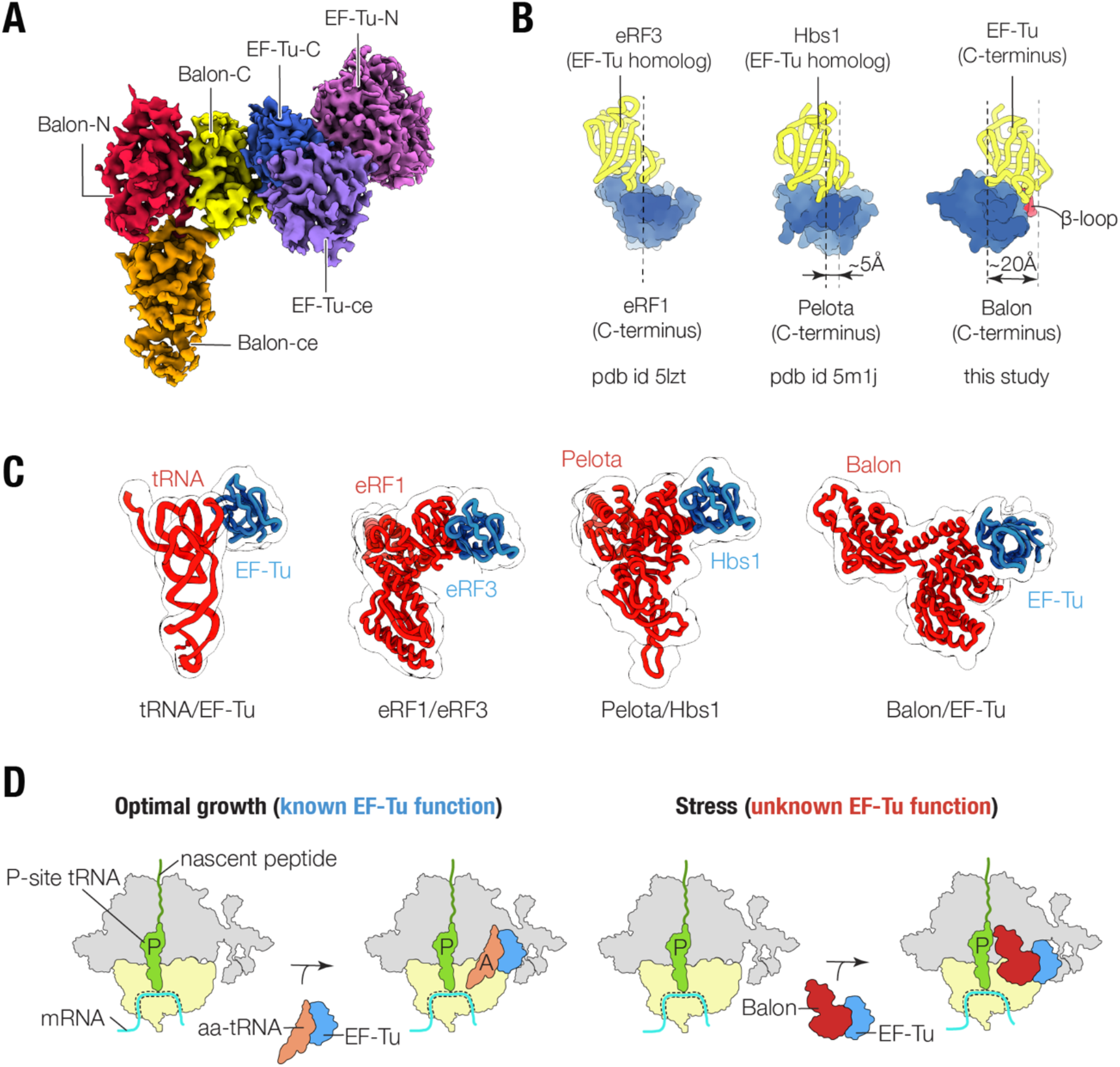
EF-Tu can act as a stress-response factor delivering Balon to the ribosomal active site. (**A**) Selected region of the cryo-EM map from the second dataset reveals the overall structure of EF-Tu bound to Balon in the ribosomal A site. (**B**) Comparison illustrating differences in binding of aRF1-type proteins (yellow) to EF-Tu-type proteins (blue), with dotted lines illustrating a shift of the protein-protein interface in Balon/EF-Tu and Pelota/Hbs1 complexes compared to the eRF1/eRF3 complex. (**C**) Side-by-side structure comparison illustrating EF-Tu-binding modes between ligands of the ribosomal A site. (**D**) Schematic representation of the new biological activity of elongation factor EF-Tu identified in this study.

### Balon occupies ribosomal active centres and multiple drug-binding sites

To understand the mechanism by which Balon recognizes idle ribosomes during the cold-shock response, we analysed its interactions with the ribosome. Balon spans two active centres of the ribosome: the decoding centre and the peptidyl-transferase centre (PTC) (**Fig. 4A-C**). Notably, as is typical for ribosome dormancy factors, the binding site for Balon overlaps with multiple binding sites of ribosome-targeting antibiotics, including tuberactinomycins^47^, oxazolidinones^48^, thermorubin^49^, and sparsomycin^50^ (**Fig. 4D-F**).

**Fig 4.**
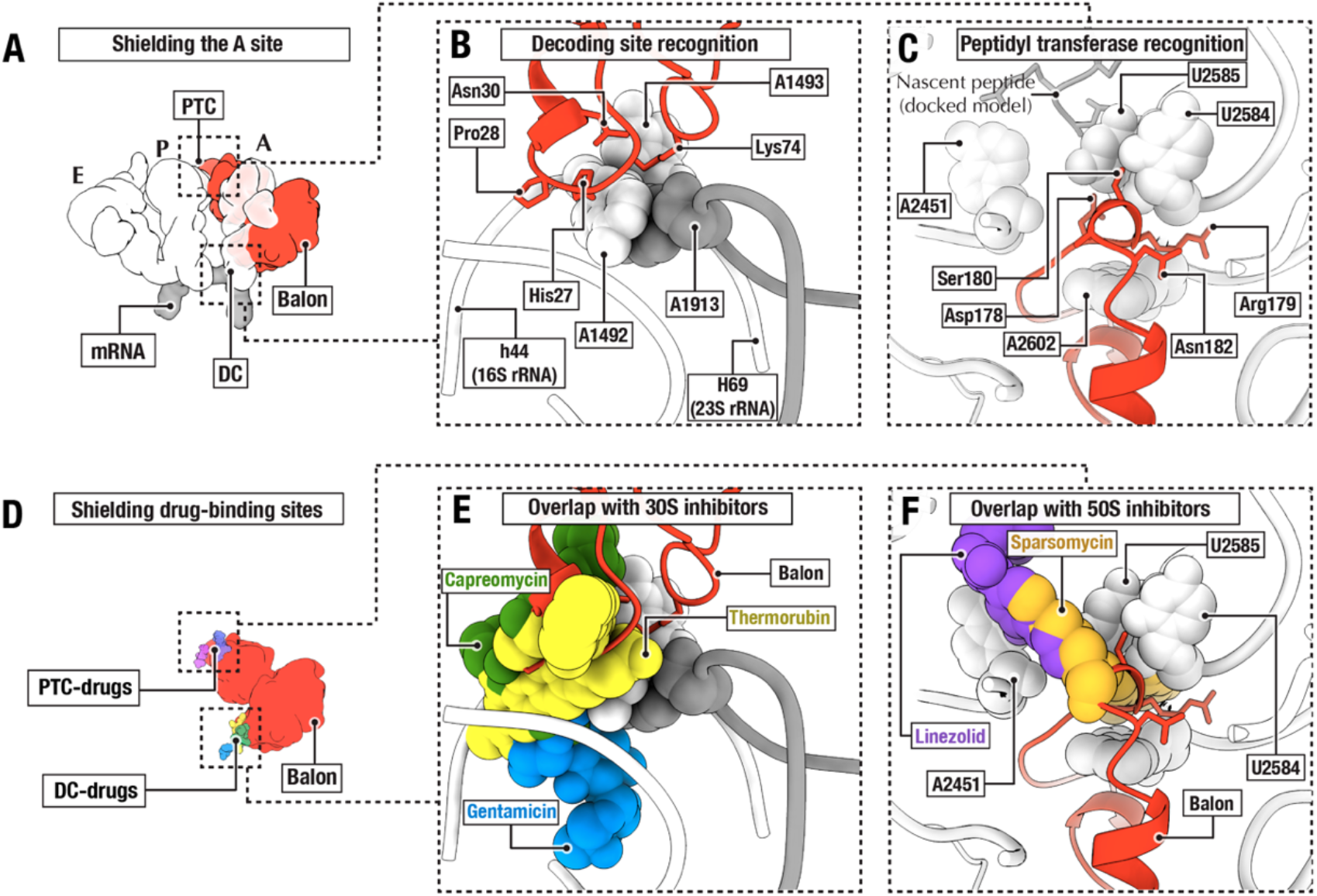
Balon occupies ribosomal active centres and overlaps with multiple drug-binding sites. (**A**) Superposition of Balon (red), tRNAs (white) and mRNA (grey) to compare ribosomal binding sites of these molecules. Balon occupies the ribosomal A site—the primary entry site of protein synthesis substrates. (**B**) Zoomed-in view of the decoding centre of the ribosome, showing details of small subunit recognition by Balon. Similar to aeRF1 and Pelota, Balon binds the ribosomal decoding centre. However, unlike aeRF1 and Pelota, Balon uses a unique HP-motif (red) which allows binding to the decoding centre without making contacts with mRNA or the mRNA entry channel. This mRNA-independent recognition strategy allows Balon to bind ribosomes concurrently with the hibernation factor RaiA. (**C**) Zoomed-in view of the peptidyl-transferase centre of the ribosome, showing details of large subunit recognition by Balon. Like aeRF1 and Pelota, Balon binds the vicinity of the peptidyl-transferase centre of the ribosome. However, unlike aeRF1 and Pelota, Balon contacts the peptidyl-transferase residue U2585 using a lasso-like protein loop (red) that wraps around the drug-binding residue A2602. (**D**) Superposition of Balon (red) and eight ribosome-targeting antibiotics (colour coded) shows that Balon occludes binding sites for drugs targeting both the peptidyl-transferase centre (PTC-drugs) and the decoding centre (DC-drugs). (**E, F**) Zoomed-in views of the decoding centre and the peptidyl-transferase centre. Balon binds multiple ribosomal drug-binding residues, overlapping with the binding sites of several families of ribosome-targeting antibiotics.

To recognize the decoding centre, Balon uses a unique strategy compared to aeRF1 and Pelota (**Fig. 4B**). Unlike aeRF1, Balon lacks the characteristic NIKS-motif necessary for mRNA stop codon recognition^51,52^. Additionally, Balon lacks the β-loop that allows Pelota to bind the vacant mRNA channel^53-55^. Instead, Balon has a distinctive HP-motif that directly binds the decoding centre at residues A1492 in helix h44, and A1913 in helix H69 (using the *E. coli* rRNA numbering) (**Fig. 4B**). These direct contacts with the decoding centre allow Balon to stay ∼ 10 Å away from the mRNA-binding site, facilitating its unique ability to bind ribosomes independently of whether mRNA is present or not. This allows Balon to bind ribosomes simultaneously with the hibernation factor RaiA, although Balon and RaiA do not form extensive contacts with each other. This concurrent binding would be impossible for aeRF1 or Pelota due to a steric clash between their N-terminal domains and the analogous position that RaiA occupies (**Fig. S8**).

We next examined the structural mechanism that Balon uses to recognize the PTC by binding 23S rRNA. Balon lacks the characteristic GGQ-motif of aeRF1 that mediates its ability to trigger nascent peptide release^56^. Instead, Balon bears a lasso-like protein loop that wraps around an rRNA nucleotide (A2602 in the 23S rRNA), positioning Balon adjacent to the outer wall of the PTC. In this position, Balon remains excluded from the catalytic centre by the 23S rRNA residue U2585, thus being unable to reach the nascent peptide (**Fig. 4C**).

### The catalytic site of Balon-bound ribosomes has an inactive conformation

Previously, the ribosomal catalytic site was observed to exist in two distinct states during protein synthesis: the inert state and the active state^57,58^. The inert (or non-induced) state was observed in ribosomes engaged in substrate search with a vacant A site, in which the 23S rRNA protects the nascent peptide to prevent spontaneous hydrolysis of the peptidyl-tRNA which would result in premature termination of protein synthesis. On the other hand, the active conformation is reached by engagement with a proper A-site substrate—such as an aminoacyl-tRNA, a translation termination factor, or an antibiotic—thereby exposing the nascent peptide for the reaction with the A-site substrate^57,58^. Our structural analysis revealed that, even though Balon binds to the ribosomal catalytic site, the structure of the catalytic site remains inert, as evident from the conformation of the 23S rRNA bases U2585 and U2506 (**Fig. 5A, B**). Balon binding to translating ribosomes therefore preserves the catalytic site in its inert state, leaving the nascent peptide inaccessible to water molecules and spontaneous hydrolysis. These findings further support the role of Balon as a *bona fide* dormancy factor and explain the well-defined electron density of the nascent peptide observed in our cryo-EM map (**Fig. S5**).

**Fig 5.**
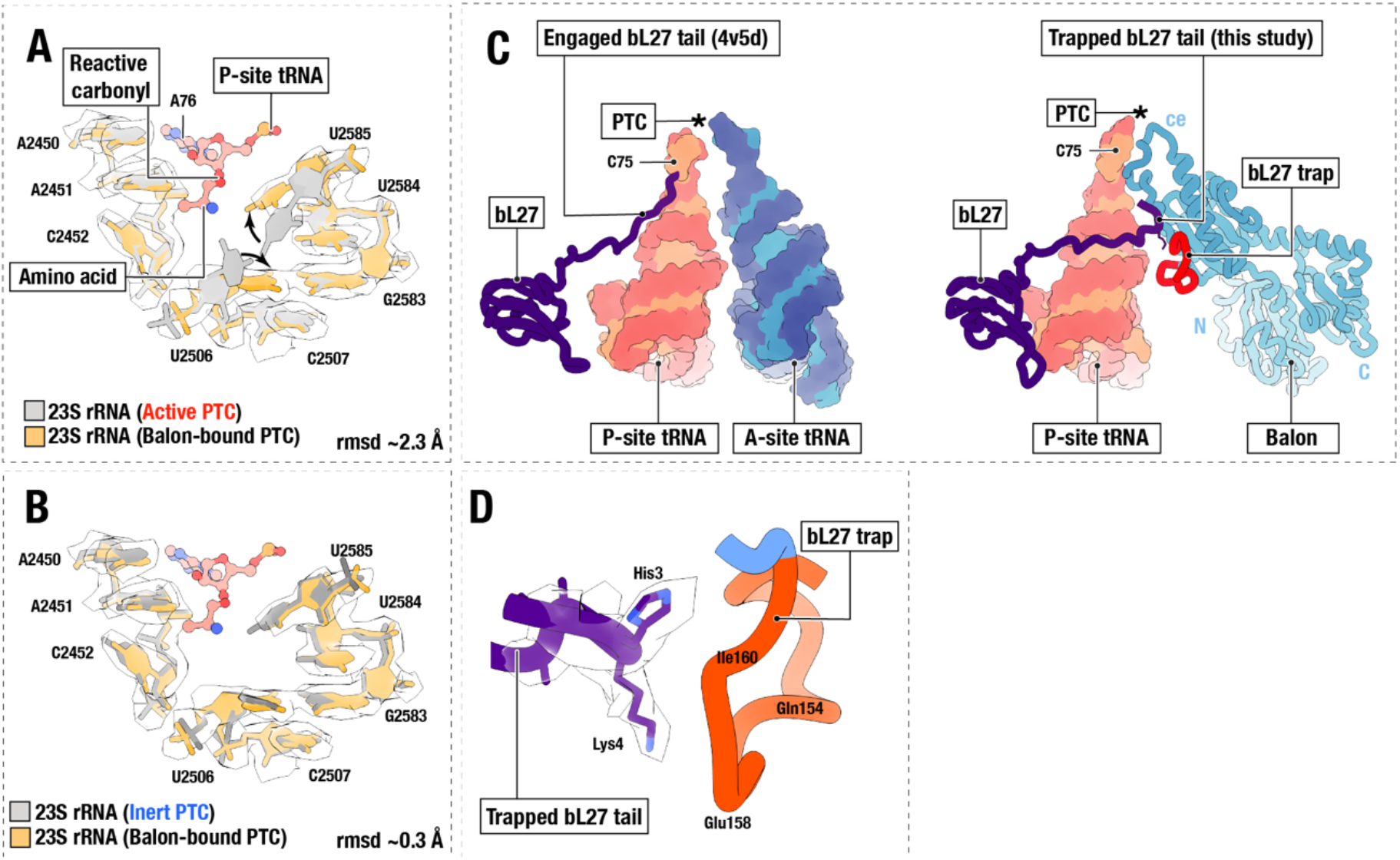
Balon-bound ribosomes have an inactive conformation of the catalytic site. (**A**) Two superposed structures compare two functional states of the ribosomal catalytic site: the active state that was observed in ribosomes just before peptide bond formation (PDB ID 4v5d) and the Balon-bound state observed in this study. The cryo-EM map shows conformation of 23S rRNA nucleotides in the Balon-bound ribosome as well as the P-site tRNA. Arrows indicate the motion of 23S rRNA residues U2585 and U2506 that would be required for the activation of the ribosomal catalytic site. (**B**) Two superposed structures compare two functional states of the ribosomal catalytic site: the inert state that was observed in ribosomes with a vacant A site (PDB ID 5mzd) and the Balon-bound state observed in this study. The rmsd values in panel (**A, B**) are shown to compare the conformation of the nucleotides U2506 and U2585 in the two states of the peptidyl-transferase center shown in each panel. (**C**) Comparison of the ribosomal protein bL27 (dark blue) in the structure of actively translating ribosomes (left) and ribosomes bound with Balon (right). (**D**) A segment of the cryo-EM map showing bL27 protein bound to bL27 trap of Balon in the structure of the 70S ribosome in complex with Balon, mRNA and P-site tRNA.

Aside from binding to 23S rRNA residues, our analysis revealed that Balon contacts another component of the bacterial peptidyl transferase centre: the ribosomal protein bL27 that is not present in eukaryotic or archaeal ribosomes. During protein synthesis in bacteria, the N-terminal tail of bL27 binds in the vicinity of the PTC and promotes ribosomal catalytic activity by positioning water molecules to favour the chemical activity of the ribosome^59,60^. However, when Balon binds bacterial ribosomes, it sequesters the N-terminal tail of bL27 using a unique loop (absent in other aeRF1-type proteins) which we term the “bL27-trap”, moving bL27 away from the peptidyl-transferase centre (**Fig. 5C, D**). Thus, unlike canonical A-site substrates, Balon binds the ribosomal A site without changing the inert structure of the ribosomal catalytic site. This indicates that Balon can preserve actively translating ribosomes in an intact state, making it theoretically possible for these ribosomes to resume translation when conditions improve.

Taken together, the overall similarity to aeRF1 and Pelota allows Balon to bind the ribosomal A site, but distinctive structural features—such as the HP-motif, the lasso-loop and the bL27-trap—appear to help Balon bind idle ribosomes in a unique fashion, independent of whether they are still engaged with mRNA and peptidyl-tRNA.

### Balon is a widespread bacterial gene that is often located in stress-response operons

We first sought to understand the distribution and conservation of Balon across all bacteria. We performed a Markov model-based homology search, using the database of reference proteomes. Notably, we observed the absence of Balon homologs in many bacteria that are commonly used to study ribosome dormancy, such as *Escherichia coli, Staphylococcus aureus* and *Bacillus subtilis*, explaining why this protein has not been previously identified. However, we were able to detect Balon homologs in 1,573 out of 8,761 representative bacteria from 23 of 27 major bacterial phyla, including many commonly studied microorganisms, such as *Thermus thermophilus*, as well as human pathogens like *Mycobacterium tuberculosis* (**Supplementary Data S3**). In all these species, Balon homologs have a similar length (360-420 amino acids) and lack the GGQ and NIKS/NIKL motifs of archaeal and eukaryotic aeRF1 family members. The Balon family members we uncovered all possess characteristic N-terminal insertions (the HP-motif) that mediate Balon’s association with the decoding centre of the ribosome. Also, all of the genomes that carry a gene for Balon also carry a gene for the hibernation factor RaiA, but almost none of them encode the hibernation factor Rmf: only 0.2% of representative bacteria simultaneously possess genes for Balon, Rmf and RaiA (**Supplementary Data S3-S9, Fig. S10-S12**). Overall, we found that Balon homologs carrying the conserved structural features detailed above are present in approximately 20 % of bacteria, always in genomes that encode RaiA homologs.

We next analysed the genomic context of Balon-coding genes. Whilst these have different genetic surroundings, they are typically located in operons that encode stress response factors, including the ribosome hibernation factor RaiA, or factors of thermal shock (e.g. Hsp20), osmotic stress (e.g. OsmC, OsmY), acid stress (e.g. HdeD), response to antibiotics (e.g. MarC, EmrB) or factors that mediate ribosome repair from nucleolytic damage (RtcB)^61-64^ (**Fig. 6C, Fig. S10**). Intriguingly, we also observed that many bacteria (603 representative species) possess multiple copies of genes for Balon, ranging from 2 to 4 copies per genome (**Fig. 6D**). For example, species *Mycobacterium* bear up to four copies of Balon-like genes, with one of these genes located in the vicinity of the hypoxia-response factor Hrp1, and another being adjacent to the gene encoding the multidrug transporter EmpB (**Fig. 6E**). Overall, this suggests that Balon is used as a stress-response protein by a wide range of bacterial species.

**Fig 6.**
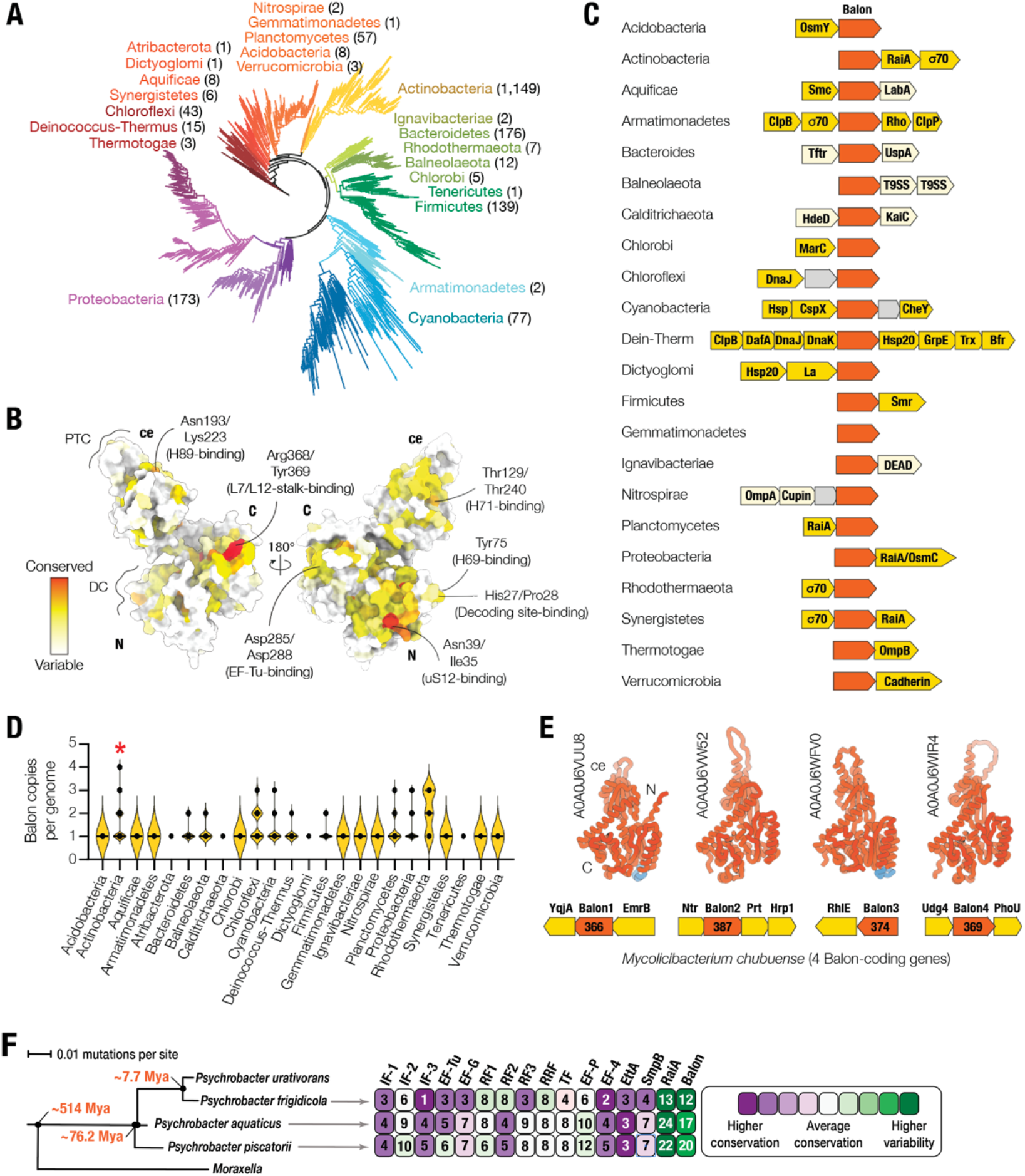
Balon-coding genes are widespread among bacteria and are often found in stress-response operons. (**A**) The bacterial tree of life shows that Balon homologs are found in most bacterial lineages (23 different phyla, encompassing 1,572 representative bacteria). (**B**) Atomic model of Balon (coloured by sequence conservation) illustrates high conservation of residues responsible for ribosome recognition. (**C**) Operon schematics illustrating the genetic context of Balon-coding genes (orange) in selected bacterial phyla. Balon-coding genes are typically found in operons that encode stress response factors (yellow). These include heat- and cold-shock response proteins (such as chaperones Hsp20, DnaK, DnaJ), alternative σ70 factors, factors of acid tolerance (HdeD) and osmotic stress tolerance (OsmB and OsmY), ribosome hibernation (RaiA), ribosome and tRNA repair (RtcB), and multidrug resistance (Smr, MarC, EmrB). (**D**) In 38% of the bacterial species analysed (603 species), Balon homologs are encoded by two, three or four gene copies located in different genomic loci, suggesting their independent expression. (**E**) As an example of a genome with a large number of Balon orthologs, here we depict the four operons encoding Balon orthologs from *Mycolicibacterium chubuense*. Notably, one of these copies (Balon 1) resides in an operon with the multidrug export protein EmrB, and another copy (Balon 2) is located in an operon with hypoxia-response factors. Their predicted structures (Alphafold) indicate a common core architecture with variations between each ortholog. (**F**) The evolutionary relationships between different *Psychrobacter* species encoding Balon orthologs, together with a comparison of the conservation of their encoded translation factors (with Balon on the far right). The numbers in each square illustrate the percentage of residues that are mutated in the indicated protein compared to its homolog in *P. urativorans*. Each square is colour-coded according to its relative level of conservation (purple = high conservation; green = low conservation) in relation to the average protein conservation between *P. urativorans* and each *Psychrobacter* species.

Notably, while searching for Balon homologs, we found that some of them have been annotated as stress-response proteins based on omics studies of *Mycobacterium* species (**Fig. S10**)^65-67^. For example, in *Mycobacterium tuberculosis* and *Mycobacterium smegmatis*, Balon homologs are annotated as “uncharacterized protein Rv2629*”* that is transcriptionally activated during cellular dormancy, leading to enhanced rifampicin resistance and increased bacterial survival and pathogenicity^65-67^. Also, in *Mycobacterium tuberculosis*, the gene for Balon is located within the same operon as RtcB, an RNA ligase that repairs the decoding site of ribosomes during starvation or stress^61-64^.

Finally, we found that Balon has an anomalously high rate of sequence evolution. In contrast to the highly conserved translation factors involved in the normal cycle of protein synthesis or quality control, Balon has a rate of molecular evolution that exceeds the average protein sequence mutation rate by approximately threefold. This places Balon in a similar category as previously identified hibernation factors such as RaiA, which exhibit similar rates of sequence evolution reflecting their relatively simple function as “molecular plugs” of vacant ribosomal active centres (**Fig. 6E**).

## Discussion

### Balon is a novel factor of bacterial ribosome dormancy

Here, by investigating an understudied psychrophilic bacterium under the conditions of cold stress, we have identified a new translation factor involved in the process of ribosome dormancy in approximately 20% of bacteria. Balon shares structural and sequence similarity with archaeal and eukaryotic translation factors, rather than the two previously described bacterial ribosome hibernation factors. It also possesses a unique ribosomal binding site and structure that allows it to bind both vacant and actively translating ribosomes: a property that sets it apart from all other known hibernation factors.

Previous studies have designated multiple bacterial, archaeal, and eukaryotic proteins as ribosome hibernation factors based on the following two criteria: their ability to bind to all or almost all ribosomes under conditions of stress, starvation, dormancy, or sporulation; and their ability to occupy the active sites of the ribosome, making ribosomes structurally incompatible with normal protein synthesis. Here we find that Balon binds and occupies the A site of at least 66 % of ribosomes in *P. urativorans* cells in response to ice treatment, thus making Balon functionally analogous to such factors as bacterial Rmf and RaiA and eukaryotic Stm1^23^, Serpbp1^19^, Irfd2^24,25^, Lso2^26-28^, and Mdf1^29,30^. RaiA and Rmf have been shown to enhance bacterial survival by protecting ribosomes from nucleolytic degradation during starvation or stress^16-18^, and it is likely that Balon functions in a similar but non-redundant manner. Our structural studies suggest that Balon protects features of the A site that remain vulnerable even in the presence of RaiA and Rmf (**Fig. S12**), providing a key missing link in our understanding of ribosome protection during translational arrest.

### Is hibernation limited to vacant ribosomes, or can translating ribosomes hibernate too?

Importantly, our discovery of Balon revises the model of ribosome dormancy and argues against a single generalized mechanism inferred from studies in *E. coli*^1,8,12^. Until now, ribosomes were believed to enter hibernation only in their vacant state: while new rounds of initiation cannot occur after dormancy-inducing stress is encountered, it was thought that elongating ribosomes would complete their existing process before entering dormancy. However, Balon has a unique property compared to RaiA and Rmf: in addition to binding vacant ribosomes, it can bind to ribosomes simultaneously bound to mRNA and peptidyl-tRNA. In our experimental cold shock conditions, about a quarter of ribosomes remain bound to mRNA and P-site tRNA while their A site is occupied with Balon, providing strong evidence that arrest of actively translating ribosomes is an important and physiologically relevant addition to the existing model.

One possible benefit of this mechanism is that ribosomes can respond to stress faster—in the middle of an elongation cycle—thereby protecting their active sites immediately without having to wait for the mRNA translation cycle to complete. We reason that this more instantaneous mode of ribosome protection may be particularly important in slow-growing bacteria with slower translation rates: where the synthesis of a single protein takes minutes rather than seconds, we envision Balon would play a critical role in pausing and protecting the substantial fraction of cellular ribosomes actively engaged in elongation when stress is encountered. This model is consistent with the recent data suggesting that ribosome hibernation factors are expressed at a basal level in both bacterial and eukaryotic cells during normal growth conditions^8^. For example, in yeast, where this process is studied in most details, cells produce of the hibernation factor Stm1 during the normal growth conditions, however Stm1 remains phosphorylated by the cellular kinase TORC1, which prevents its binding to ribosomes. During starvation and stress, Stm1 becomes de-phosphorylated, leading to its association with the ribosome^68^.

One future question to address will be: what happens to Balon-bound ribosomes when conditions improve? One possible scenario is that Balon dissociates from the A site, and ribosomes that were bound to mRNA and peptidyl-tRNA resume translation. Alternatively, these ribosomes may undergo recycling, in which bound mRNA and peptidyl-tRNA would be degraded before a new translation cycle can begin. Therefore, it will be exiting to elucidate the fitness benefits of Balon-mediated ribosome dormancy in different stress conditions and in organisms with different translation rates.

Notably, our finding that Balon can associate with tRNA-bound ribosomes has important implications for the current strategies to identify hibernating ribosomes using cryo-EM, which rely on elimination of ribosomes bound to P-site tRNA at the early stages of particle classification. Our study strongly suggests that this approach can lead to an incomplete or even misleading understanding of ribosome hibernation by eliminating a significant fraction of cellular ribosomes that may also be hibernating. Our work suggests that a more effective approach to identifying hibernating ribosomes should involve inspecting all active sites of the ribosome for the presence of stress-response factors.

### EF-Tu as a factor of stress response

It is well-established that EF-Tu plays an essential role in the normal protein synthesis cycle by delivering aminoacyl-tRNAs to the ribosomal A site^69^. Our work, however, identifies a new role for EF-Tu in the response to stress. Similar to EF-Tu homologs that deliver aeRF1 and Pelota to ribosomes in archaea and eukaryotes^46,54^, EF-Tu appears to deliver Balon to ribosomes in bacteria. Whilst the relative affinities of EF-Tu for Balon and aminoacyl tRNA are unknown, the observation of EF-Tu bound to Balon in our cryo-EM maps implies that they are at least somewhat comparable. Various stress and starvation conditions are known to decrease the availability of aminoacylated tRNAs, which are the primary binding partners of EF-Tu^70-72^. Therefore, it is plausible that, under these conditions, greater amounts of free EF-Tu would become available for association with Balon, facilitating its delivery to ribosomes. In this way, Balon recruitment may be regulated by the ratio of uncharged/charged tRNAs within a cell, providing a rapid response to starvation. This model is consistent with our mass-spectrometry data illustrating that our ice treatment causes only a minor increase in the intracellular levels of both Balon and RaiA, implying that actively growing *P. urativorans* cells possess a sufficiently large pool of Balon to protect ribosomes immediately, without having to rely on transcription and translation of stress response operons when the availability of nucleotides and amino acids may be limiting. Regardless of the exact mechanism of Balon delivery, our discovery of EF-Tu binding to Balon is significant because EF-Tu is the target of several antibiotics (e.g. GE2270A and kirromycin) and understanding its mechanism of action serves as the basis for developing safer and more effective clinical drugs^73-75^.

### Balon provides evidence that bacteria possess functional eukaryote-type translation factors

Comparative analyses of bacterial genomes have predicted dozens of hypothetical homologs of eukaryotic and archaeal protein synthesis factors in bacteria. These apparent homologs were initially identified for eukaryotic translation initiation factors, including eIF2, eIF3, eIF4A and eIF6^76^. More recently, bioinformatics efforts have identified homologs of archaeo-eukaryotic translation factors aeRF1 and Pelota in various bacterial genomes, suggesting a much higher degree of compositional diversity in the bacterial translation apparatus than previously thought^34,35^. However, many of these genes are highly variable, fragmented and poorly conserved, in defiance of expectations for core translation machinery^76,77^. In the absence of functional and structural data, it has remained unclear whether these apparent homologs function in bacterial translation, or even whether their protein products are produced.

Our work establishes a role for Balon in ribosome dormancy and provides strong evidence for structural conservation with the aeRF1-family of translation factors. The sporadic presence of Balon-coding genes across the bacterial tree of life suggests that these genes may have originated from aeRF1-coding genes through a horizontal gene transfer from archaea or eukaryotes to bacteria. Our structures provide insight into the replacement of the archaeo-eukaryotic GGQ- and NIKS-motifs with the HP-motif, lasso-loop, and bL27-trap. These features allow Balon to uniquely engage with the decoding site without making any contacts with the mRNA channel. This proposed evolutionary transformation appears to endow Balon with the ability to bind multiple functional states of the ribosome as opposed to one specific state that is recognized by aeRF1 or Pelota.

Irrespective of the exact evolutionary origins of Balon, our work demonstrates that bacteria and eukaryotes share a higher similarity in their translation machinery than suggested by conventional models of protein synthesis. Overall, this finding supports the idea that several previously predicted bacterial homologs of archaeo-eukaryotic translation factors may indeed participate in protein synthesis. Extrapolating beyond Balon to this larger pool of putative factors, the bacterial kingdom may possess nearly twice as many regulators of protein synthesis as currently estimated – a rich area for future investigation.

## METHODS

### Production of *P. urativorans* biomass

As a model organism, we used the bacterium *Psychrobacter urativorans*. Freeze-dried cells of *P. urativorans* were obtained from the American Type Culture Collection (ATCC 15174). The cell pellet was rehydrated in 15 mL of pre-chilled Marine Broth 2216 media (Sigma-Aldrich) and incubated in a shaker (SciQuip Incu-Shake Mini) at 150 rpm at 10 °C for 7 days, according to the ATCC protocol. This culture was then used to inoculate 1L of pre-chilled Marine Broth 2216 media and incubated at 150 rpm for 4 days at 10 °C until the culture reached an OD_600_ of 0.272. The cells were then transferred on ice for 10 min and centrifuged for 5 min at 4 °C and 5,000 g, yielding approximately 1 g of cell pellet.

### Ribosome isolation

To lyse the cells, the pellet was rapidly resuspended in 1 mL of buffer A (50mM Tris-HCl pH 7.5, 20 mM Mg(OAc)_2_, and 50 mM KCl), transferred to 2 mL microcentrifuge tubes containing approximately 0.1 mL of 0.5 mm zirconium beads (Sigma-Aldrich BeadBug^™^), and disrupted by shaking for 30 sec at 6.5 m/s speed in a bead beater (Thermo FastPrep FP120 Cell Disrupter). The sample was then centrifuged for 5 min at 4 °C and 16,000 g to remove cell debris, and the resulting supernatant was collected and centrifuged for 1 min at 16,000 rpm and 4 °C to remove the remaining debris. To analyse polysome profiles, we analysed 0.1 mL of crude *P. urativorans* lysates per time point, using 10-40 % sucrose gradients in buffer A after 3 hours of centrifugation at 35,000 rpm and 4 °C in SW41 rotor (Beckman Coulter). To isolate ribosomes for structural analysis, the cell lysate corresponding to 30 min of ice treatment was then mixed with PEG 20,000 (25 % w/v) to a final concentration of 0.5 % (w/v) and centrifuged for 5 min at 4 °C and 16,000 g to precipitate insoluble aggregates. Then, the supernatant was mixed with PEG 20,000 (powder) to the final concentration of ∼12.5 % (w/v) and centrifuged for 5 min at 4 °C and 16,000 g to precipitate ribosomes. To monitor precipitation of ribosomes, we analysed lysates and their fractions using size-exclusion chromatography with Superdex Increase 200 10/300 in buffer A (**Fig. S1**). The obtained ribosome-containing pellet was dissolved in 50 μL of buffer A, and the solution was passed twice through PD Spin Trap G-25 microspin columns (GE Healthcare) to clear crude ribosomes from small molecules. The obtained solution had an OD_260_ of 34.89 and OD_260/280_ of 1.71, corresponding to a ribosome concentration of 512 nM. This solution was split into 10 μL aliquots and frozen at -20 °C for subsequent cryo-EM and mass spectrometry analyses.

### Mass-spectrometry analysis of crude samples of *P. urativorans* ribosomes

For each measurement shown in **Supplementary Data S1 and S2**, a 10 μL aliquot of crude *P. urativorans* ribosomes solution was reduced with 4.5 mM dithiothreitol and heated at 55 °C. The sample was alkylated with the addition of 10 mM iodoacetamide before proteolytic digestion with 0.2 μg Promega sequencing grade trypsin and incubation at 37 °C for 16 h. The resulting peptides were desalted by Millipore C18 ZipTip, following the manufacturer’s protocol, with final elution into aqueous 50 % (v/v) acetonitrile. Desalted peptides were dried under vacuum before being resuspended in aqueous 0.1 % trifluoroacetic acid (v/v) for LC-MS/MS.

Peptides were loaded onto a mClass nanoflow UPLC system (Waters) equipped with a nanoEaze M/Z Symmetry 100 Å C18, 5 µm trap column (180 µm x 20 mm, Waters) and a PepMap, 2 µm, 100 Å, C18 EasyNano nanocapillary column (75 μm x 500 mm, Thermo). The trap wash solvent was aqueous 0.05% (v/v) trifluoroacetic acid and the trapping flow rate was 15 µL/min. The trap was washed for 5 min before switching flow to the capillary column. Separation used gradient elution of two solvents: solvent A, aqueous 0.1 % (v/v) formic acid; solvent B, acetonitrile containing 0.1 % (v/v) formic acid. The flow rate for the capillary column was 330 nL/min and the column temperature was 40 °C. The linear multi-step gradient profile was: 3-10 % B over 7 mins, 10-35 % B over 30 mins, 35-99 % B over 5 mins and then proceeded to wash with 99 % solvent B for 4 min. The column was returned to initial conditions and re-equilibrated for 15 min before subsequent injections.

The nanoLC system was interfaced with an Orbitrap Fusion Tribrid mass spectrometer (Thermo) with an EasyNano ionization source (Thermo). Positive ESI-MS and MS2 spectra were acquired using Xcalibur software (version 4.0, Thermo). Instrument source settings were: ion spray voltage—1,900 V; sweep gas— 0 Arb; ion transfer tube temperature—275 °C. MS1 spectra were acquired in the Orbitrap with 120,000 resolution, the scan range of m/z 375-1,500; the AGC target of 4e5, and the max fill time of 100 ms. Data dependent acquisition was performed in top speed mode using a 1 s cycle, selecting the most intense precursors with charge states >1. Easy-IC was used for internal calibration. Dynamic exclusion was performed for 50 s post precursor selection and a minimum threshold for fragmentation was set at 5e3. MS2 spectra were acquired in the linear ion trap with: scan rate—turbo; quadrupole isolation—1.6 m/z; activation type—HCD; activation energy—32%; AGC target—5e3; first mass—110 m/z; max fill time—100 ms. Acquisitions were arranged by Xcalibur to inject ions for all available parallelizable time.

Peak lists in Thermo .raw format were converted to .mgf using MSConvert (version 3.0, ProteoWizard) before submitting to database searching against the *P. urativorans* subset of the UniProt database (3^rd^ Aug 2022, 2,349 sequences; 769,448 residues)^78^ appended with 118 common proteomic contaminants. Mascot Daemon (version 2.6.0, Matrix Science) was used to submit the search to a locally running copy of the Mascot program (Matrix Science Ltd., version 2.7.0). Search criteria specified: enzyme—trypsin; max missed cleavages—2; fixed modifications—carbamidomethylation of protein C-termini; variable modifications—acetylation of protein N-termini, deamidation of Asn and Gln residues, N-terminal conversion of Gln and Glu to pyro-Glu, oxidation of Met, and phosphorylation of Ser, Thr and Tyr residues; peptide tolerance—3 ppm; MS/MS tolerance—0.5 Da; instrument—ESI-TRAP. Peptide identifications were passed through the percolator algorithm to achieve a 1% false discovery rate assessed against a reverse database. The search data are summarized in (**Supplementary Data S1 and S2**), where molar percentages of each identified protein were calculated from Mascot emPAI values by expressing individual values as a percentage of the sum of all emPAI values in the sample, according to Ref.^79^. To calculate the relative abundance of each cellular protein before and after 30 minutes of ice treatment (as shown in **Supplementary Data S1**), their total spectrum counts in the ice treated sample were divided by the corresponding total spectrum counts of the control (non-ice-treated) sample. An infinite value for a few proteins means that in the control sample we have not been able to detect evidence for a protein by spectral counting.

### Cryo-EM grid preparation and data collection

To prepare ribosome samples for cryo-EM analyses, 10 μL aliquots of crude ribosomes were thawed on ice and loaded onto glow discharged (20 mA, 90s, PELCO easiGlow) Quantifoil grids (R1.2/1.3, 200 mesh, copper), using 2 μL of the sample per grid. The grids were then blotted for 1 or 2 sec at 100% humidity (using blotting force -5) and vitrified using liquid nitrogen-cooled ethane in a Vitrobot Mark IV (Thermo Fisher Scientific). The grids were then used for data collection with a 200 kV Glacios electron cryo-microscope (Thermo Scientific) with Falcon 4 detector located at the York Structural Biology Laboratory, University of York, UK. For each movie, the grids were exposed to a total dose of 78 e/Å2 across 4.95 seconds. A nominal magnification of 240,000 x was applied, resulting in a final calibrated object sampling of 0.574 Å pixel size. 8,903 micrograph movies were recorded in aberration free image shift (AFIS) mode using a defocus targets of -1.25, -1.0, -0.75, -0.5 μm.

### Cryo-EM data processing

Cryo-EM data for the first *P. urativorans* dataset were processed using RELION 3.1^80^ as summarised in **Table S1, Fig. S2, Fig. S4**). In brief, a total of 180,467 particles were picked from 8,903 micrographs using the Laplacian of Gaussian picker (250 – 300 Å particle diameter; 0.75 stdev threshold). Particle images were initially downscaled three-fold and extracted in a 280 pixel box (1.722 Å effective pixel size). Two rounds of 2D classification were carried out to clean the dataset, with 83,896 ‘good’ particles selected for 3D refinement. This generated an initial map at 3.7 Å resolution, in which heterogeneity was apparent at the Balon binding site and decoding centre. To resolve this, angular assignments from this refinement job were used to carry out a masked classification without alignment, focusing on the Balon density and the P-site. This effectively separated particles into three groups corresponding to differential factor occupancy: 1—Ribosome(empty); 2—Ribosome/Balon/RaiA; and 3— Ribosome/Balon/tRNA/mRNA. The final subsets of Ribosome/Balon/RaiA and Ribosome/Balon/tRNA/mRNA particles were re-extracted in a 420 pixel box (1.148 Å effective pixel size) and subjected to per-particle CTF refinement, polishing and post-processing. Finally, sharpened maps weighted by estimated local resolution were calculated. All reported estimates of resolution are based on the gold standard Fourier shell correlation (FSC) at 0.143, and the calculated FSC is derived from comparisons between reconstructions from two independently refined half-sets.

Very weak density was apparent for EF-Tu in both of these reconstructions, indicative of very low occupancy. Following further screening, a second dataset was collected from another grid (**Table S1**). This was processed using cryoSPARC v3.3.266^81^ as shown in **Fig. S3, Fig. S4.**

In brief, after patch motion correction and CTF estimation, particles were picked from 3,706 micrographs using cryoSPARC blob picker (180 – 220 Å particle diameter). Particles were then extracted using a box size of 700 pixels and subjected to two rounds of 2D classification. The resulting classes (62,815 particles) were selected for ab-initio reconstruction and homogenous refinement with C1 symmetry, which resulted in the map with a resolution of 3.36 Å. This map (Ribosome/Balon/EF-Tu) has better density for EF-Tu but is heterogeneous with respect to both RaiA and tRNA in the P-site. Further focused classification as above demonstrated that EF-Tu is associated with both Ribosome/Balon/RaiA and Ribosome/Balon/tRNA/mRNA particles in the second dataset.

### Model building, refinement and deposition

The atomic models of *P. urativorans* ribosomes and the ribosome-binding proteins were produced using Coot v0.8.9.2^82^ and AlphaFold^83^. As a starting model, we used the atomic model of ribosomal proteins generated by AlphaFold and the atomic model of rRNA from the coordinates of *Thermus thermophilus* ribosomes (PDB ID 4y4o)^84^. These rRNA and protein models were morph-fitted into the cryo-EM maps using ChimeraX^85^ and Phenix^86^ and then rebuilt using Coot based on the information about the genomic sequence of *P. urativorans* (RefSeq GCF_001298525.1). In the ribosome complex with Balon, mRNA and tRNA, the mRNA molecule was modelled as poly-U, and the tRNA molecule was modelled as U_1-72_A_73_C_74_C_75_A_76_.

The density corresponding to Balon was initially identified as a non-ribosomal protein, which was initially modeled as a poly-alanine chain to determine its backbone structure. This poly-alanine backbone model was then used as an input file for a search of proteins with similar fold in the Protein Data Bank using the NCBI tool for tracking structural similarities of macromolecules, Vast^87^. This search identified the archaeal protein aeRF1 from *Aeropyrum pernix* as the most similar known structure to Balon, suggesting that Balon is a bacterial homolog of aeRF1 (**Table S2**). We therefore searched for *P. urativorans* proteins that have a similar sequence to that of *A. pernix* aeRF1. Using three iterations of Markov model-based search with HHMER^88^ in the UniProtKB database, we found that *P. urativorans* encodes a hypothetical protein (Uniport id A0A0M3V8U3) with sequence similarity to aeRF1 and Pelota. This protein, which we termed Balon, had a sequence that matched the cryo-EM map and was used to create its atomic model. The resulting atomic structures of Balon in complex with the ribosome, RaiA and EF-Tu or Balon in complex with the ribosome, tRNA and mRNA were then refined using Phenix real space refinement, and the refined coordinates were validated using MolProbity^89^ and awaiting to be deposited in the Protein Data Bank. To compare corresponding domains in the structures of Balon, aeRF1 and Pelota, we used a flexible secondary structure-based alignment with FatCat^90^.

### Evolutionary analysis of Balon

To assess phylogenetic distribution of Balon in bacterial species, we performed three iterations of homology search using the sequence of Balon from *P. urativorans* (Uniprot id A0A0M3V8U3) as an input for a profile hidden Markov Model-based analysis with HMMER^88^. For each search iteration, we used the following search options: -E 1 --domE 1 --incE 0.01 --incdomE 0.03 --seqdb uniprotrefprot. The resulting dataset was reduced first by removing protein sequences that lacked information about their Phylum (21 sequences), then by removing sequences that were shorter than 300 amino acids as they typically lacked one or two of the three domains of Balon/aeRF1 (which included 806 sequences), then by removing sequences that were annotated as a protein fragment (34 sequences), and finally by removing duplicated sequences (31 sequences). This resulted in the dataset including 1,896 sequences of Balon homologs from 1,565 bacterial species (**Supplementary Data 2**).

To gain insight into a possible evolutionary origin of Balon from the archaeo-eukaryotic family of aeRF1 proteins, we performed a complementary analysis in which we searched for bacterial homologs of the archaeal aRF1 using three iterations of HMMER^88^. As an input for the first iteration, we used the sequence of aRF1 from the archaeon *A. pernix* (Uniprot id Q9YAF1), which we identified as being one of the closest structural homologs of Balon. For each iteration, we used the database of Reference Proteomes restricted to the bacterial domain of life, using these search options: -E 1 --domE 1 --incE 0.01 --incdomE 0.03 --seqdb uniprotrefprot. The resulting dataset was reduced first by removing sequences lacking information about their Phylum (21 sequences), then by removing sequences which were lacking at least one of the three domains of aeRF1 proteins (sequences shorter than 300 amino acids, which included 1,422 sequences), then by removing sequences annotated as a protein fragment (5 sequences), and finally by removing duplicated sequences (104 sequences). This resulted in the dataset of 1,617 sequences of bacterial aeRF1 homologs from 1,353 bacterial species (**Supplementary Data 3**).

To map the identified Balon homologs on the tree of life, we combined the results of the previous searches in the (**Supplementary Data 2**) and (**Supplementary Data 3**) and removed repetitive entries, which resulted in a dataset of 1,898 protein sequences from 1,572 bacterial species (**Supplementary Data 4**). We then aligned the combined sequences using Clustal Omega^91^ with default parameters, which resulted in a multiple sequence alignment (Supplementary Data 5) and a phylogenetic tree (**Supplementary Data 6**), which were analysed with JalView^92^ and iTol^93^, respectively. To compare phylogenetic distribution of Balon, Rmf and RaiA-type hibernation factors, we have repeated homology search using HMMER for Rmf (using the *E. coli* sequence of Rmf as an input) (**Supplementary Data 7**) and RaiA (using the *E. coli* sequences of RaiA as an input) (**Supplementary Data 8**). To compare rates of sequence evolution in Balon and other translation factors, we used Uniport align/pairwise comparison tool to calculate pairwise similarities and TimeTree5^94^ to estimate an approximate age of *Psychrobacter* species.

## Acknowledgements

We thank Babak Javid (UCSF Centre for Tuberculosis) and Zofia Lightowlers, Bert van der Berg and Heath Murray (Newcastle University) for their thoughtful comments on the manuscript, and Dr Johan Turkenburg and Sam Hart for their work supporting the York cryo-EM facility. This work was funded by the Newcastle University NUORS 2021 Award (to K.H-B.), the Medical Research Council (MR/N013840/1 to C.L.E.), the BBSRC UK (BB/T008695/1. to C.R.B.) a UKRI Future Leader Fellowship (MR/T040742/1 to J.B.), a Wellcome Trust & Royal Society Sir Henry Dale Fellowship (221818/Z/20/Z to C.H.H.), and the Royal Society (RGS/R2/202003 to S.V.M.). This project was undertaken on the NSBL Cluster and the Viking Cluster, which are high-performance compute facilities provided by Newcastle University and the University of York, respectively. We are grateful for computational support from the University of York High Performance Computing service, Viking, and the Research Computing team, and support from the Newcastle University Structural Biology Laboratory. We also acknowledge the York cryo-EM facility supported by Wellcome Trust (206161/Z/17/Z) and the York Centre of Excellence in Mass Spectrometry that was created with a capital investment through Science City York and supported by EPSRC (EP/K039660/1; EP/M028127/1) and Yorkshire Forward with funds from the Northern Way Initiative. For the purpose of open access, the authors have applied a CC BY public copyright license to any Author Accepted Manuscript version arising from this submission.

