## Supplementary Information for "A bacterial ribosome hibernation factor with evolutionary connections to eukaryotic protein synthesis"

**Suppl. Information**

***A bacterial ribosome hibernation factor***

***To whom correspondence should be addressed**

**Tables**

Table S1 | Statistics of cryo-EM data collection and model building

Table S2 | Structurally similar proteins to Balon (using Vast)

**Figures**

Figure S1 | Ribosome isolation from the cold-adapted bacterium *P. urativorans*

Figure S2 | Cryo-EM data processing workflow using RELION 3.1

Figure S3 | Cryo-EM data processing workflow using cryoSPARC

Figure S4 | Validation of the cryo-EM maps

Figure S5 | Cryo-EM maps of ribosome ligands in Balon-bound ribosomes

Figure S6 | Balon is a tRNA-mimicking protein that binds the ribosome in complex with EF-Tu

Figure S7 | aeRF1 and Balon have similar folds but share just 10% of sequence identity

Figure S8 | Balon and aeRF1 have dissimilar orientations of their N-terminal domains, leading to an entirely dissimilar strategy of the decoding centre recognition

Figure S9 | Balon binding site has multiple conformations during the ribosomal active cycle

Figure S10 | Omics studies of the Balon homolog in the human pathogen *Mycobacterium tuberculosis*

Figure S11 | In the *P. urativorans* genome, Balon and RaiA genes reside in different operons

Figure S12 | Relative binding sites and distribution of Balon, Rmf and RaiA in bacterial species

**Supplementary Data**

Supplementary Data 1 | Mass-spectrometry analysis of ice-treated vs control *P. urativorans* samples

Supplementary Data 2 | Mass-spectrometry analysis of crude *P. urativorans* ribosomes

Supplementary Data 3 | aeRF1 homologs in bacterial species (by HMMER)

Supplementary Data 4 | Balon homologs in bacterial species (by HMMER)

Supplementary Data 5 | Combined Balon homologs in bacterial species

(combined Suppl. Data 3 and 4) (by HMMER)

Supplementary Data 6 | Aligned sequences of Balon homologs in bacteria (by ClustalOmega)

Supplementary Data 7 | Tree of life calculated using Balon sequences (by ClustalOmega)

Supplementary Data 8 | RaiA homologs in bacterial species (by HMMER)

Supplementary Data 9 | Rmf homologs in bacterial species (by HMMER)

**Maps and Models**

<https://drive.google.com/drive/folders/1ynWIWq7N6pWEGcdhkdusa2s8BXr5ntA6?usp=share_link>

**Table S1 | Cryo-EM data collection, refinement and validation statistics.**

|  | 70S/Balon/RaiA  (RELION, EMBD-xxx)  (PDB xxxx) | 70S/Balon/tRNA/mRNA  (RELION, EMBD-xxx)  (PDB xxxx) | 70S/Balon/EF-Tu  (cryoSPARC, EMBD-xxx) (PDB xxxx) |
| --- | --- | --- | --- |
| **Data collection and processing** |  |  |  |
| Magnification | 240k | 240k | 150k |
| Voltage (kV) | 200 kV | 200 kV | 200 kV |
| Electron exposure (e–/Å^2^) | 78 | 78 | 50 |
| Defocus range (μm) | -0.5,-0.75,-1.0,-1.25 | -0.5,-0.75,-1.0,-1.25 | -0.5,-0.75,-1.0,-1.25 |
| Pixel size (Å) | 0.574 | 0.574 | 0.934 |
| Symmetry imposed | C1 | C1 | C1 |
| Final particle images (no.) | 31,356 | 23,994 | 62,815 |
| Map resolution (Å)  FSC threshold | 3.1  0.143 | 3.1  0.143 | 3.36  0.143 |
| Map resolution range (Å) | 2.87 – 50.0 | 2.85 – 50.0 | 3.36 - 53.3 |
| **Refinement** |  |  |  |
| Model resolution (Å)  FSC threshold | 3.1(masked)  3.1 (unmasked)  0.5 | 3.1(masked)  3.1 (unmasked)  0.5 | 3.4(masked)  3.5 (unmasked)  0.5 |
| Model resolution range (Å) | 3.1-50 | 3.1-50 | 3.36-50 |
| Map sharpening *B* factor (Å^2^) | -55.7 | -56.8 | -57.5 |
| Model composition  Non-hydrogen atoms  Protein residues  Ligands | 142,411  6,225  1 (Mg) | 143,467  6,120  1 (Mg) | 142,623  6,331  1 (Mg) |
| *B* factors (Å^2^) (min/max/mean)  Protein  Nucleotide  Ligand | 3.65/60.39/31.03  0.78/99.22/31.15  6.77/6.77/6.77 | 3.65/60.39/31.00  0.78/119.49/32.06  6.77/6.77/6.77 | 10.21/90.45/37.67 10.52/154.64/41.46 20.35/20.35/20.35 |
| R.m.s. deviations  Bond lengths (Å)  Bond angles (°) | 0.010  0.962 | 0.021  1.174 | 0.030  1.059 |
| Validation  MolProbity score  Clashscore  Poor rotamers (%) | 1.94  11.25  0.18 | 2.05  14.79  0.18 | 1.98  13.58  0.84 |
| Ramachandran plot  Favoured (%)  Allowed (%)  Disallowed (%) | 94.47  5.40  0.13 | 94.45  5.40  0.15 | 95.80  4.10  0.07 |

**Table S2 | Proteins that are structurally similar to Balon (identified by VAST).** The table illustrates that, of all structurally characterized proteins, Balon is the most structurally similar to proteins Pelota and aRF1 from the archaeo-eukaryotic branch of life.

| **PDB ID** | **Balon segment** | [**Aligned**](https://www.ncbi.nlm.nih.gov/Structure/VAST/vasthelp.html#VASTTable)  **residues** | **P-value** | **RMSD**  **(Å)** | **Sequence identity (%)** | **Protein** | **Organism** |
| --- | --- | --- | --- | --- | --- | --- | --- |
| [3OBY](https://www.ncbi.nlm.nih.gov/Structure/mmdb/mmdbsrv.cgi?uid=84588&Dopt=s) | All structure | 193 | 10e-6.2 | 4.0 | 15.5 | Pelota | *Archaeoglobus fulgidus* |
| [3WXM](https://www.ncbi.nlm.nih.gov/Structure/mmdb/mmdbsrv.cgi?uid=122819&Dopt=s) | Ce-domain | 117 | 0.0004 | 3.7 | 12.8 | Pelota | *Aeropyrum pernix* |
| [3IR9](https://www.ncbi.nlm.nih.gov/Structure/mmdb/mmdbsrv.cgi?uid=76511&Dopt=s) | All structure | 104 | 0.0023 | 2.6 | 12.5 | aRF1 | *Methanosarcina mazei* |
| [2QI2](https://www.ncbi.nlm.nih.gov/Structure/mmdb/mmdbsrv.cgi?uid=59472&Dopt=s) | Ce-domain | 101 | 10e-5.4 | 3.0 | 12.9 | Pelota | *Thermoplasma acidophilum* |

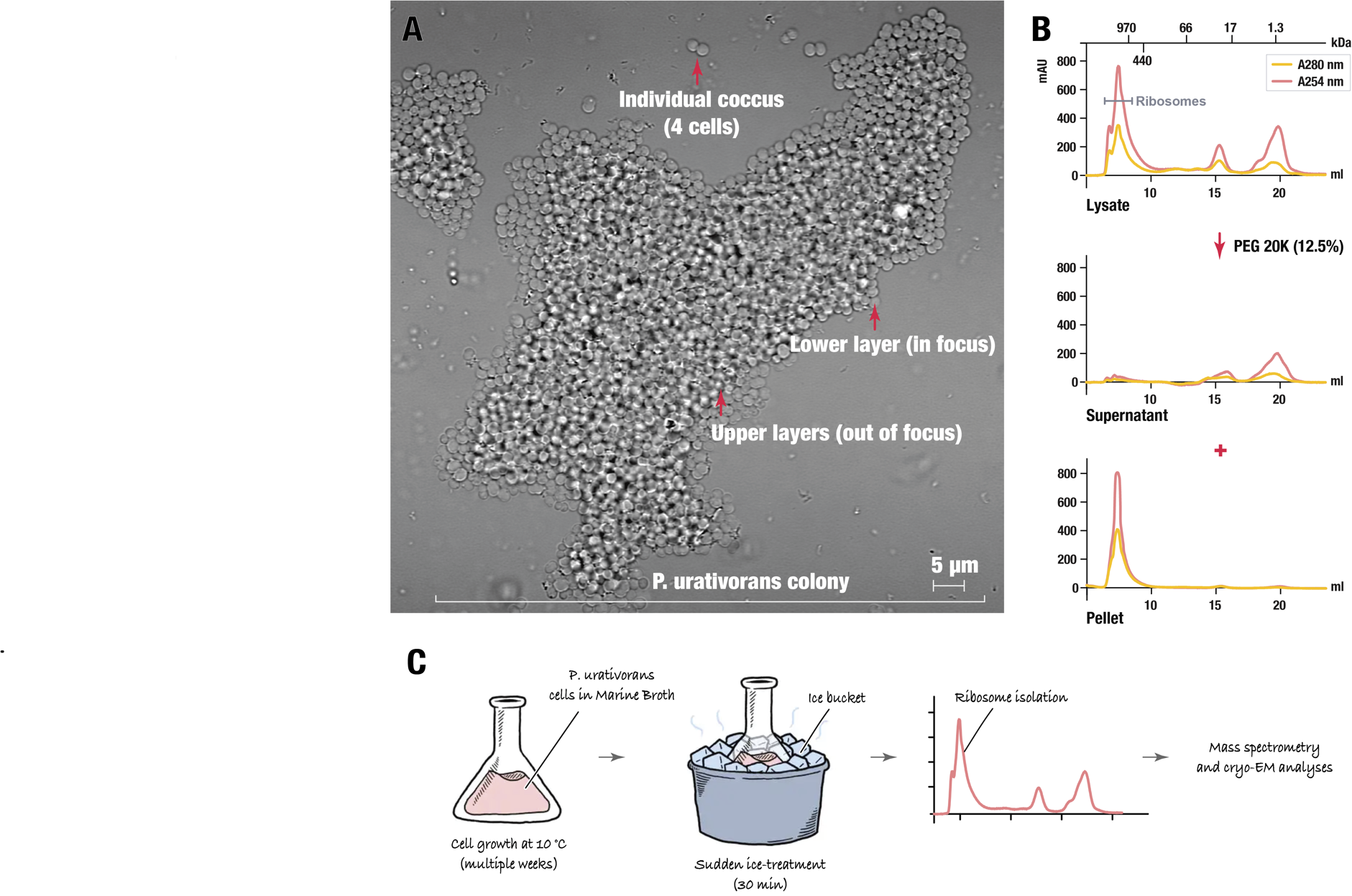

**Figure S1 | Ribosome isolation from the cold-adapted bacterium *P. urativorans***. (**A**) A brightfield microscopy image of *P. urativorans* cells used in this study shows a colony comprising approximately 1,000 cells. This colony was isolated from an actively growing liquid culture of *P. urativorans* (OD600 ~ 0.2) by transferring 10 μL of a cell suspension onto an agar bed. Unlike many common model bacteria, such as *E. coli*, *P. urativorans* is not a unicellular organism but rather a multicellular organism. Each individual cell of this bacterium is organized in a so-called coccus, which contains two, four, or more cells that are surrounded by a thick cell wall and internally divided by strongly developed cross-walls. Most of these cocci further self-assemble into larger cellular aggregates, like the one shown here. These aggregates typically comprise a few dozen to a few hundred cells arranged into carpet-like monolayers, with multiple monolayers attached to each other. This morphology, along with the presence of bright pigments in *P. urativorans* cells, makes this species unsuitable for fluorescence microscopy or cytometry studies. (**B**) Size-exclusion chromatography profiles illustrate the lysate fractionation strategy used in this study. Before PEG20K fractionation, the lysate contains particles of various sizes before the fractionation (the upper panel). However, once the PEG20K is added to the lysate, this causes selective precipitation of large particles, including ribosomes, as evident from their disappearance from the soluble fraction (the middle fraction) and their accumulation in the pellet (the lower fraction). Thus, by precipitating the content of cell lysates with PEG20K (12.5%), we were able to achieve nearly complete isolation of *P. urativorans* ribosomes from cell lysates.

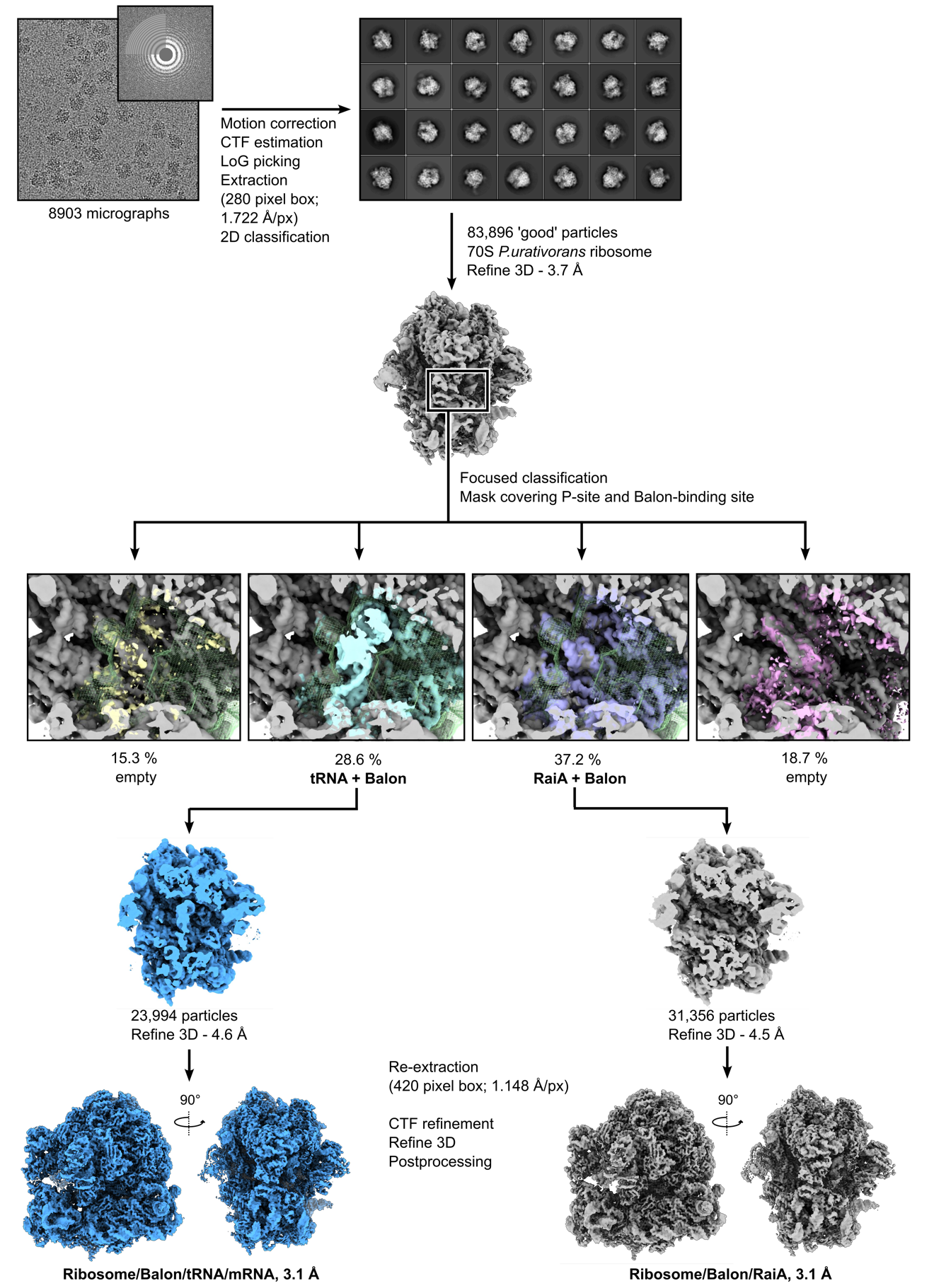

**Figure S2 | Cryo-EM data processing workflow (Dataset 1) for the reconstruction of *P. urativorans* ribosomes and focused classification analysis of Balon-bound ribosomes.** The pipeline shows a representative micrograph at 240kx with fourier transform, 2D classes, 3D reconstructions and major steps of data processing using RELION 3.1.

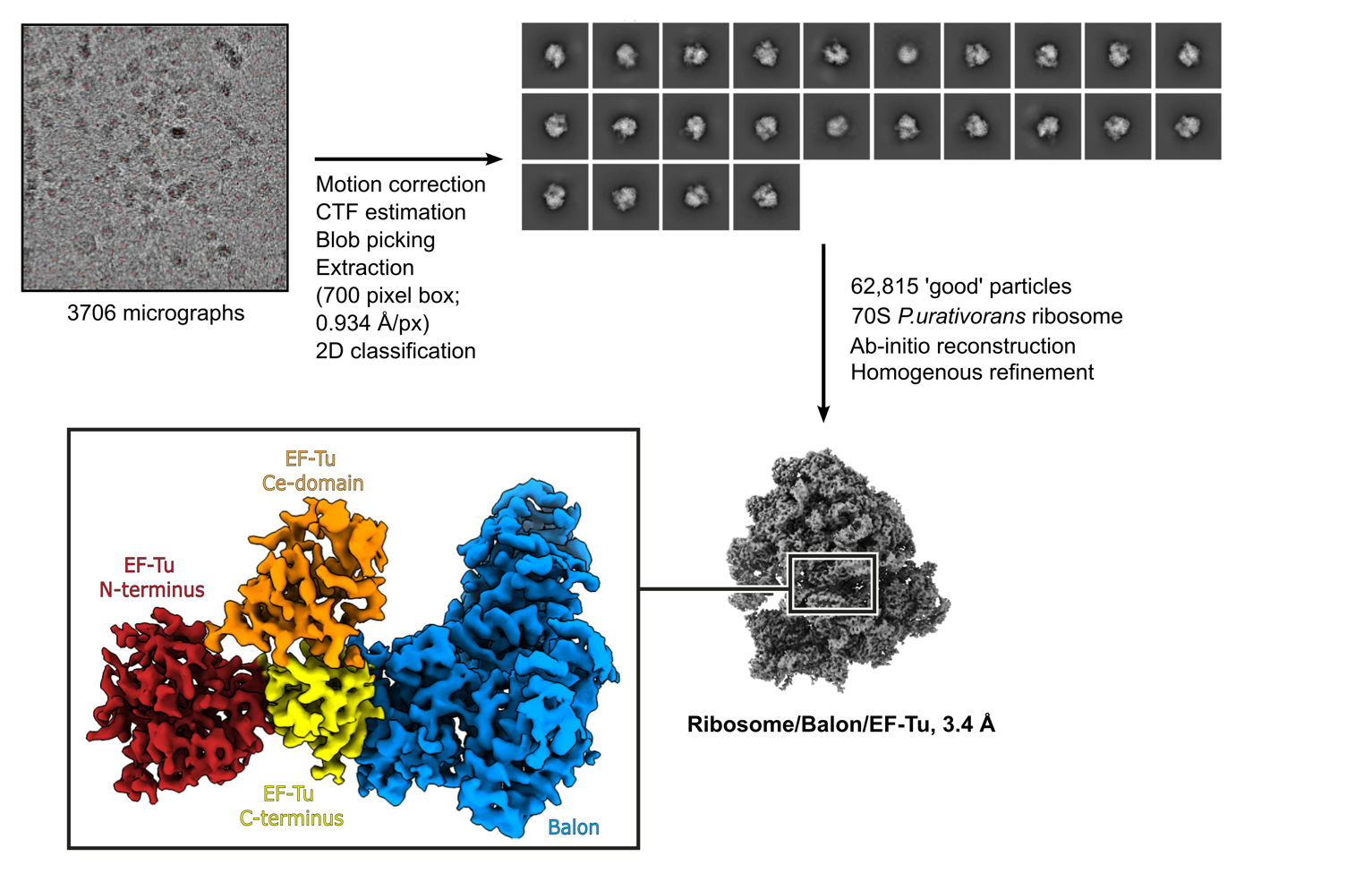

**Figure S3 | Cryo-EM data processing workflow (Dataset 2) for the reconstruction of *P. urativorans* ribosomes bound to Balon and elongation factor EF-Tu.** The pipeline shows a representative micrograph, 2D classes and major steps of data processing using cryoSPARC v.3.3.2, as well as a segment of the final cryo-EM map showing density for Balon and EF-Tu.

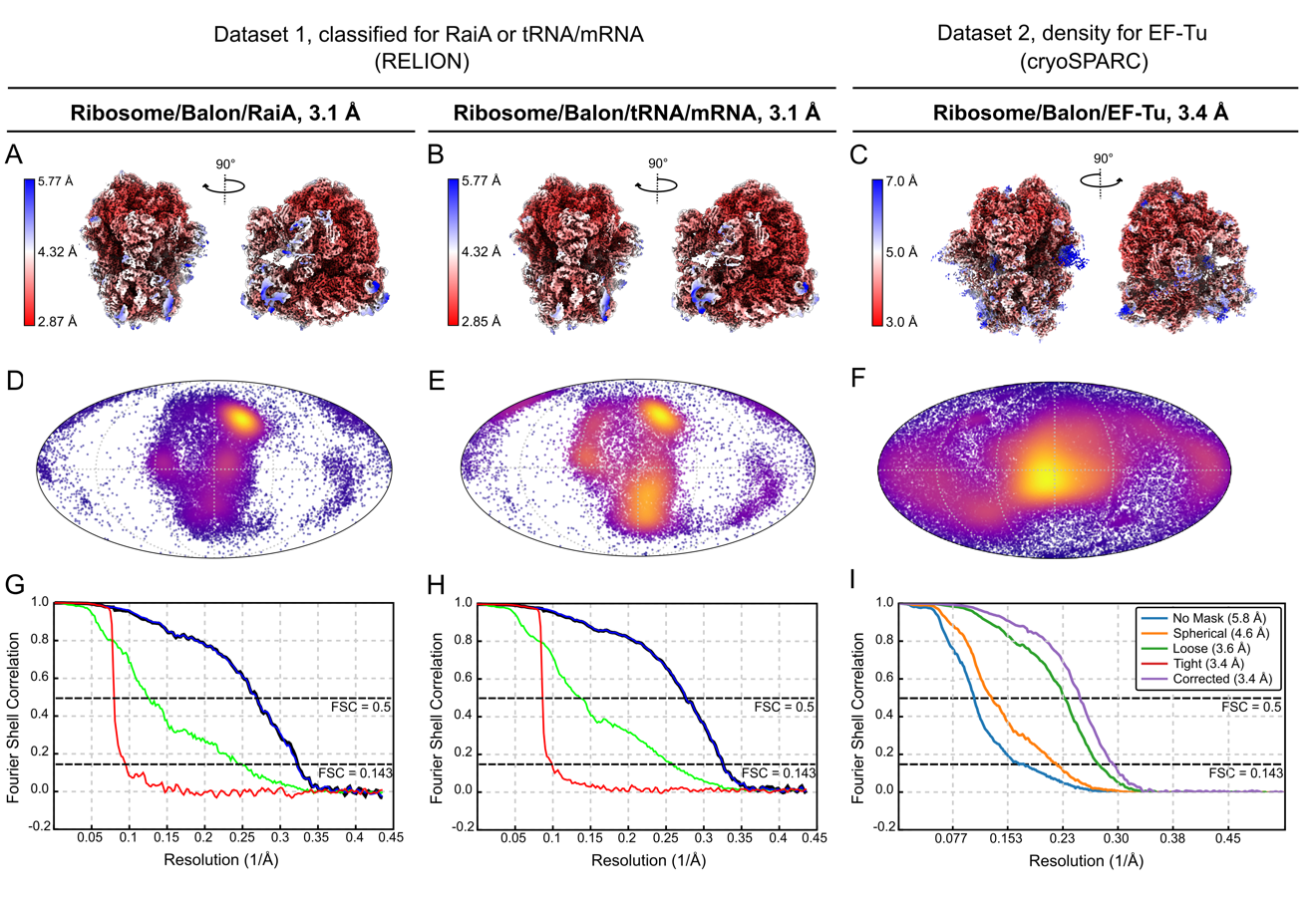

**Figure S4 | Validation of cryo-EM maps of *P. urativorans* ribosome.** Panels descriptions refer to the Ribosome/Balon/RaiA map (left), Ribosome/Balon/tRNA/mRNA map (centre) and Ribosome/Balon/EF-Tu map (right) as indicated at the top of the figure. (**A, B, C**) Final cryo-EM maps, surface coloured by estimated local resolution as indicated in the heatmap key. Two orthogonal views are shown. (**D, E, F**) Angular distribution plot of particles in the final reconstructions, shown as a Mollweide projection (**G, H**) Gold-standard Fourier shell correlation (FSC) curves for final maps generated by RELION postprocessing. Masked (blue), unmasked (green) and phase-randomised masked (red) plots are shown. (**I**) Fourier shell correlation (FSC) curves for the Ribosome/Balon/EF-Tu map, generated by cryoSPARC. Curves are labelled according to legend.

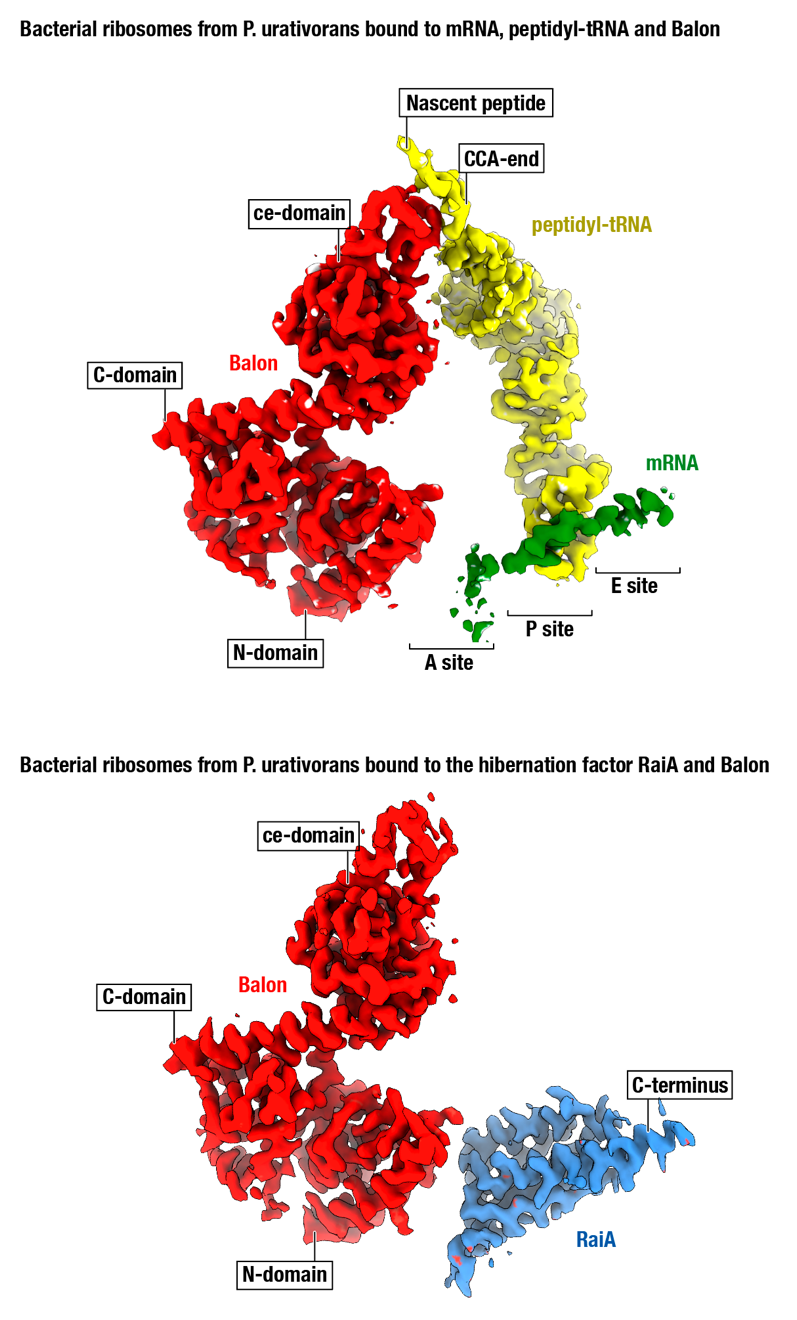

**Figure S5 | Cryo-EM density for molecules that bind ribosomes concurrently with Balon.** The cryo-EM maps show ribosomal ligands in two distinct classes of ribosomes in our dataset. The first class represents the ribosome complex that contains heterologous mRNA, heterologous peptidyl-tRNA, and Balon. The second class represents the ribosome complex that comprises Balon and the hibernation factor RaiA. In both of these complexes, Balon has a nearly identical density, illustrating its invariant binding to these two states of the ribosome.

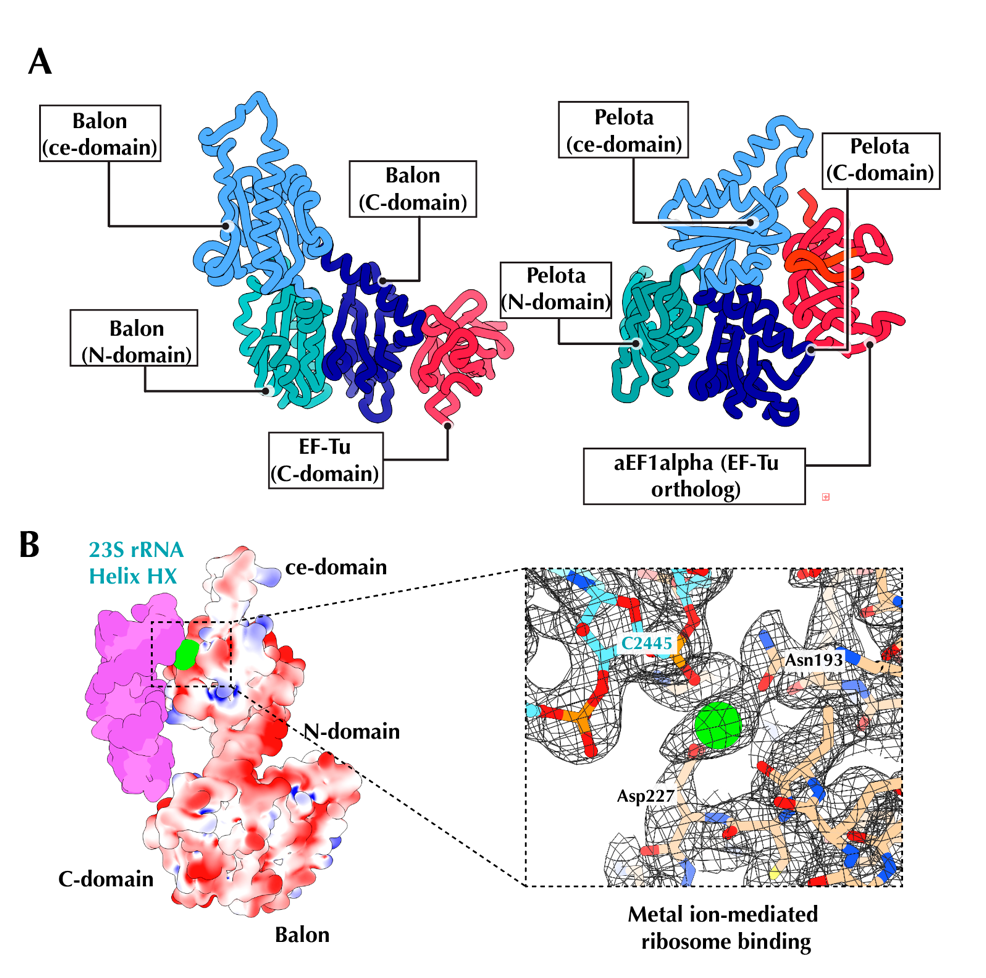

**Figure S6 | Balon mimics tRNA molecules in binding the ribosome and EF-Tu.** (**A**) Side-by-side comparison of interactions between EF-Tu (red; known as EF1a in eukaryotes and archaea) and Balon (left) or Pelota (right), each colored by domain. Both Balon and Pelota use their C-terminal domain (purple) to bind the C-terminal domain of EF-Tu/aEFa-alpha. However, in contrast to Pelota, Balon interacts with the C-terminal domain of EF-Tu using a different face of its C-terminal domain, illustrating the evolutionary divergence of EF-Tu recognition between Pelota and Balon. (**B**) The electrostatic potential surface, shown within a range between -2 (red) and +2 (blue) kT/e-, illustrates that Balon is a negatively charged protein that binds the ribosome not only via direct molecular contacts but also via contacts mediated by a metal ion.

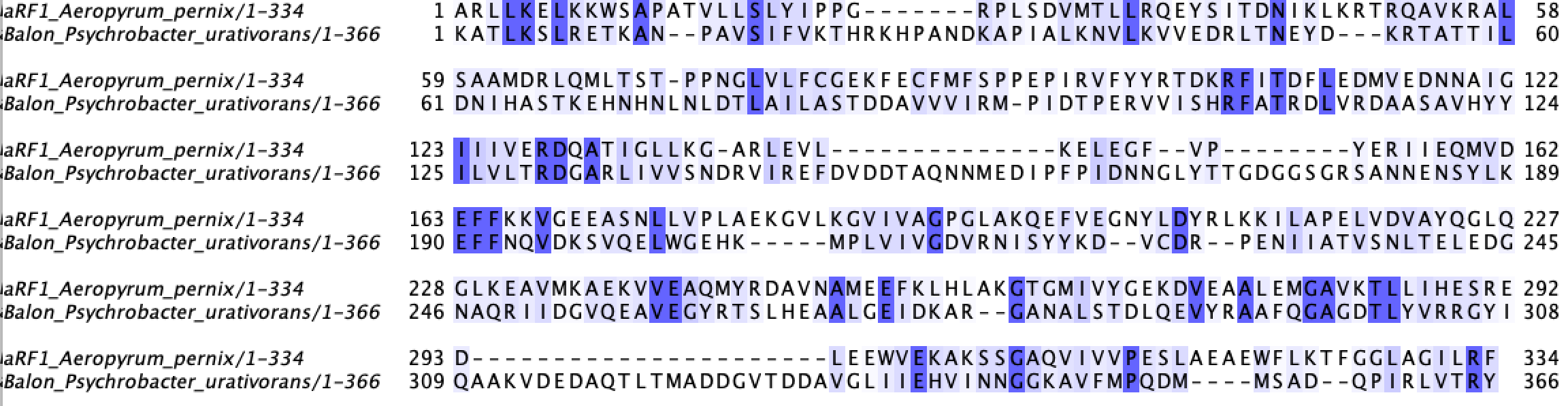

**Figure S7 | Balon and aRF1 have the same fold but share just 10% of identical residues.** The structure-guided alignment compares protein sequences of Balon from *P. urativorans* (Uniprot id A0A0M3V8U3) and the archaeal translation termination factor aRF1 from *A. pernix* (Uniprot id Q9YAF1). The sequences are aligned using an alignment of structures of Balon and aRF1 with FatCat. Balon and aRF1 have approximately **29%** sequence similarity and **10.3%** sequence identity.

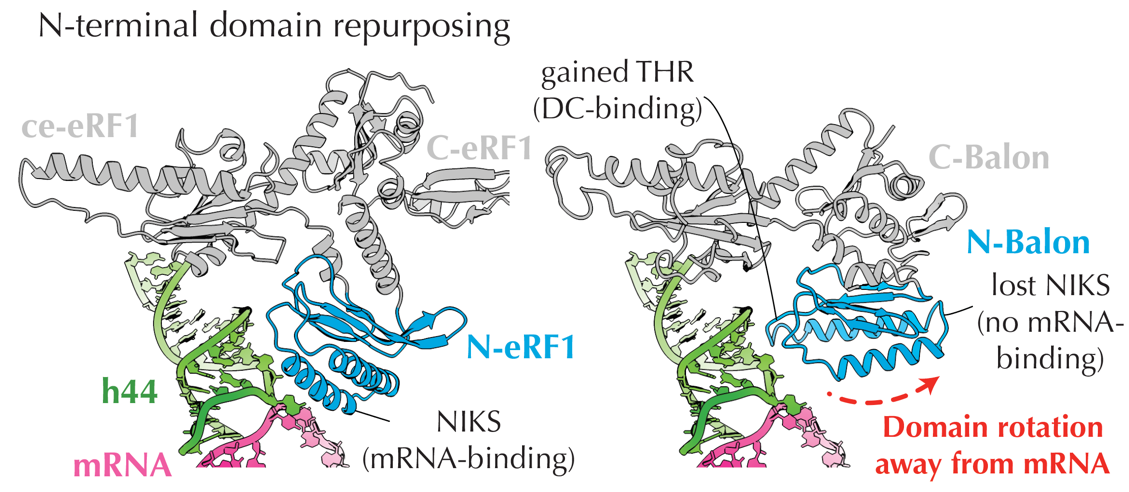

**Figure S8 | Balon and aeRF1 have similar folds but bind the ribosome using the opposing faces of their N-terminal domain.** Comparison of eRF1 (left) and Balon (right) binding to the decoding centre of the ribosome illustrates one key difference between these two factors. Despite Balon and aeRF1 proteins having the same domain organisation, the Balon N-terminal domain (blue) is rotated by about 60 degrees compared to aeRF1. As a result, Balon and aeRF1 use different surfaces of their N-terminal domains to recognize the decoding centre of the ribosome. This rotation also moves the N-terminal domain away from the mRNA-binding channel, leading to mRNA-independent recognition of the ribosome by Balon. This mRNA-independent recognition allows Balon to bind the decoding centre concurrently with RaiA (aeRF1 cannot bind concurrently with RaiA due to a clash in the mRNA channel).

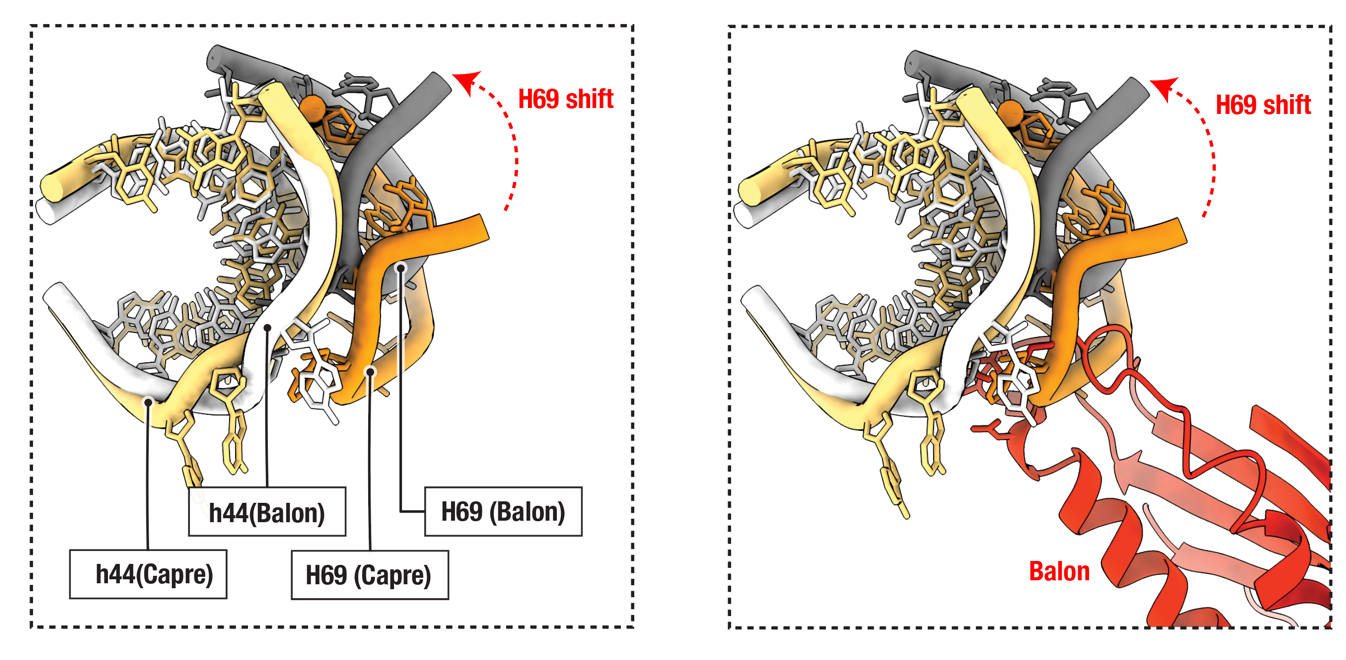

**Figure S9 |The Balon-binding site on the ribosome adopts multiple conformations during translation.** To recognize the decoding centre of the ribosome, Balon binds at the interface of two rRNA helices: 16S helix h44 and 23S helix H69. These two helices are known to adapt various conformations relative to each other. The figure shows superposition of two structures in which ribosomes adopt either a non-rotated conformation (as observed in this study) or a rotated conformation (e.g. PDB 4v7m). In the rotated state of the ribosome, H69 is shifted relative to h44, and H69 would clash with the N-terminal domain of Balon, indicating that Balon cannot use the same recognition strategy to bind this conformation of the decoding centre.

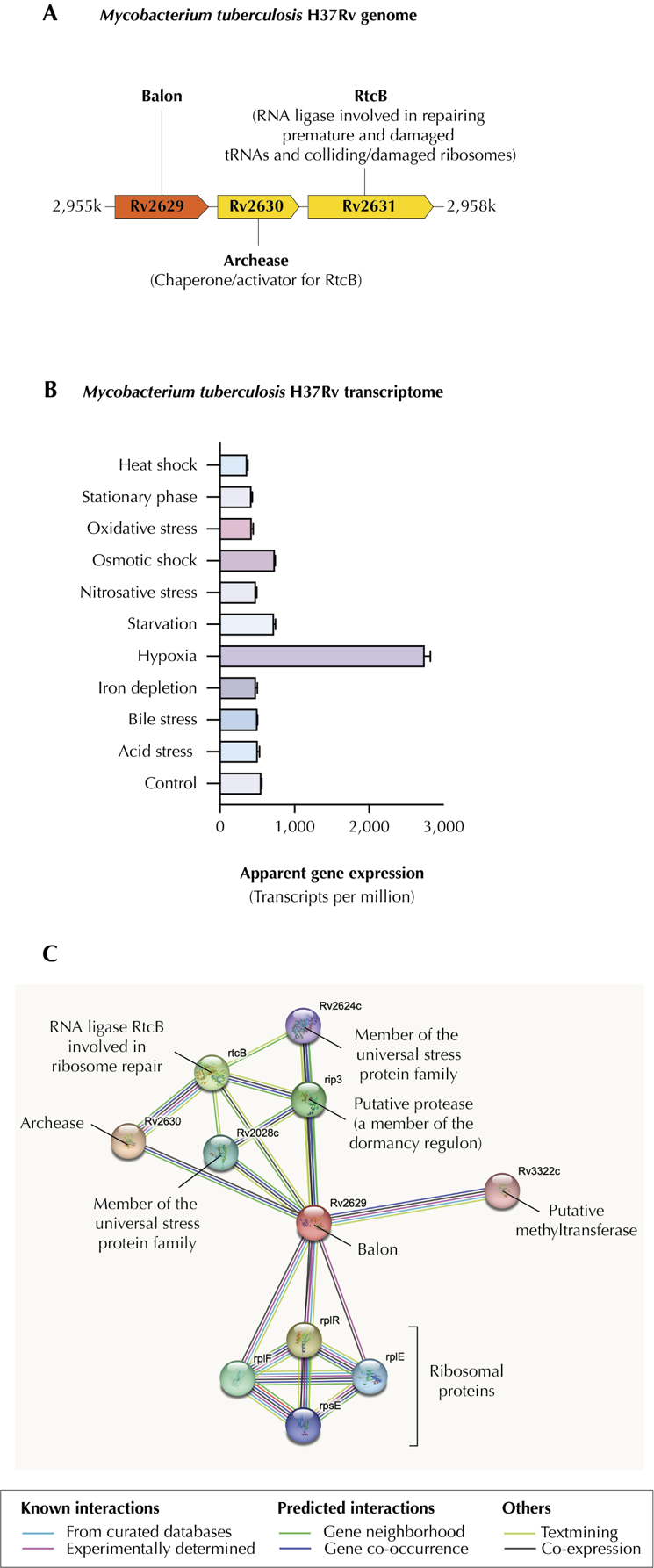

**Figure S10 | Omics studies of the Balon homolog in the human pathogen *Mycobacterium tuberculosis* H37Rv**. (**A**) Genomic surroundings of the Balon-coding gene (according to [http://tbdb.bu.edu](http://tbdb.bu.edu/), Balon ID is Rv2629). (**B**) mRNA transcription levels showing that Balon expression is activated by hypoxia and, to a lesser extent, osmotic shock and starvation (according to [www.pathogenex.org](http://www.pathogenex.org), Balon ID is WP_003413610.1). (**C**) Experimentally determined and inferred Balon interactions with other *M. tuberculosis* proteins ([www.string-db.org](http://www.string-db.org), Balon ID is Rv2629).

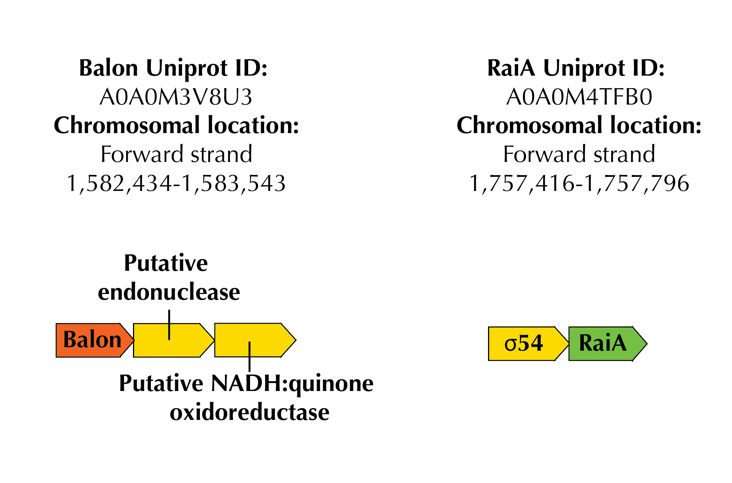

**Figure S11 | In the *P. urativorans* genome, Balon and RaiA genes reside in different operons.** In many proteobacteria, Balon- and RaiA-coding genes are adjacent to each other, consistent with their simultaneous binding of Balon and RaiA to the ribosome. In *P. urativorans*, however, they reside in different genomic loci. This example shows that simultaneous binding of these two proteins to dormant ribosomes during the cold shock does not require their genomic colocalization.

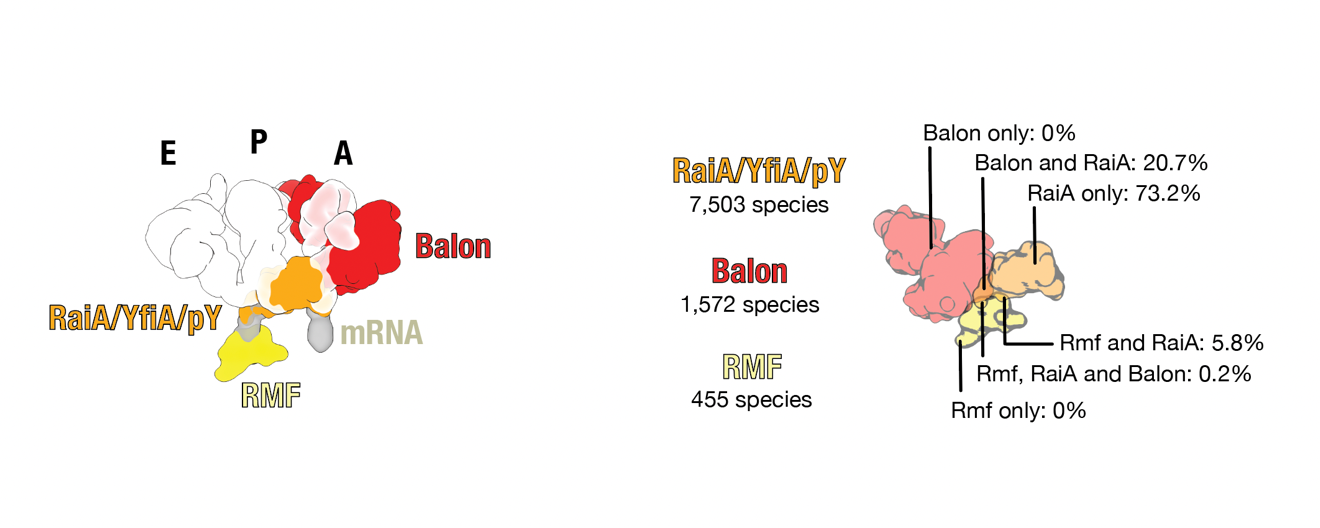

**Figure S12 | Relative binding sites and distribution of Balon, Rmf, and RaiA in bacterial species.** The left panel shows superposed structures of Balon and previously characterized bacterial hibernation factors, including RaiA-type factors and Rmf, illustrating how the ribosomal binding sites of these proteins are positioned relative to the binding sites of mRNA (grey) and tRNA (white) molecules. Because Balon possesses a structurally altered N-terminal domain (compared to aeRF1 proteins), it can bind the ribosome concurrently with the hibernation factor RaiA. This binding is impossible for aeRF1 due to the clash between RaiA and the mRNA-binding domain of aeRF1. Overall, the figure illustrates that Balon, RaiA and RMF have non-overlapping binding sites in the ribosome, making their concurrent binding possible. The right panel depicts the relative occurrence of Balon, Rmf and RaiA in representative bacterial proteomes.
